# H3K4me3 methyltransferase KMT2F promotes pre-initiation complex formation by RNA Polymerase I to regulate ribosomal RNA transcription

**DOI:** 10.1101/2024.11.04.621718

**Authors:** Kaisar Ahmad Lone, Amit Mahendra Karole, Geethanjali Ravindran, Shweta Tyagi

## Abstract

Trimethylation of histone 3 lysine 4 (H3K4me3) is mark of active transcription and its regulatory role in RNA polymerase II-mediated transcription has been well-studied. However, if and how this mark regulates RNA polymerase I (RNA Pol I) is not known. Here we used customized genome assemblies for rDNA to demonstrate that KMT2A and KMT2F bind to entire rDNA loci. The binding of these enzymes were mirrored by the binding of H3K4me2 and H3K4me3 marks. Using biochemical assays, we demonstrate the interaction of KMT2- specific subunits with RNA Pol I transcriptional machinery. Our findings reveal KMT2F as the primary KMT depositing the H3K4me3 on rDNA. Loss of H3K4me3 adversely affects the epigenetic landscape and promotes heterochromatization of rDNA locus. Mechanistically, we show that KMT2F promotes pre-initiation complex formation of RNA Pol I. Our findings highlights the thus far undiscovered role of H3K4me3 in the transcriptional initiation of rDNA genes.

## Introduction

Within the cell, the intricate coordination of cellular growth and proliferation occurs through a precisely regulated series of steps called ribosome biogenesis. Deregulation within this elaborate process is associated with a diverse spectrum of human diseases including cancer (Narla & Ebert, 2010)(Dixon et al., 2007)(Ganapathi et al., 2007)(Hannan et al., 2013) (Heiss et al. 1998). The control of ribosomal RNA (rRNA) transcription plays a central role in the complex pathway of ribosome biogenesis. The initial and crucial regulatory stage in ribosome biogenesis occurs through the transcription of ribosomal RNA genes (rDNA) by RNA polymerase I (RNA Pol I) within the dynamic membrane-less structure called nucleolus (Stults et al., 2008)(Hannan et al., 2013). The average diploid human genome harbours approximately 400 tandemly arranged repeats of rRNA genes, distributed across five acrocentric chromosomes (Stults et al., 2008). Each Mammalian transcription unit (43 kb), consists of a coding region spanning 13.3 kb interspaced by 30 kb intergenic spacer (IGS). The coding region is responsible for encoding the 47S pre-ribosomal RNA (pre-rRNA), which undergoes subsequent processing to yield the mature 5.8S, 18S, and 28S rRNA. The coding unit is flanked by the non-coding regulatory elements, promoter, and a terminator sequence in the IGS. In addition to human rDNA core or 47S promoter, the rDNA promoter contains spacer promoter which is separated by enhancer repeats and harbours essential signal sequences for transcription initiation and termination (Zentner et al., 2011)(I. Grummt, 1982)(Sollner-Webb et al., 1983)(L. Grummt et al., 1986)(A. Kuhn &Grummt, 1987); (Labhart and Reeder 1986). The main rRNA promoter (referred to as 47S promoter from hereon) directs the pre-rRNA synthesis, while the role of spacer promoter is not clear.

The activity of RNA polymerase I is intricately regulated through interactions with numerous auxiliary factors. These factors play a crucial role in promoter recognition and contribute significantly to the processes of transcription initiation, elongation, and termination (Sharifi & Bierhoff, 2018)(Goodfellow & Zomerdijk, 2013). One of the primary regulators of rDNA transcription is the upstream binding factor (UBF). It plays an important role in various stages of this process, including pre-initiation complex (PIC) assembly, promoter escape, and elongation (Panov et al., 2006)(Stefanovsky et al., 2006). Additional elements within the RNA polymerase I machinery, such as the Selectivity Factor 1 (SL1) complex and RNA polymerase I-specific transcription initiation factor RRN3, are crucial for facilitating the initiation of RNA polymerase I. The 300 kd SL1 complex, composed of the TATA binding protein (TBP) and four transcription-associated factors (TAFs) TAF1A/TAF_I_48, TAF1B/TAF_I_63, TAF1C/TAF_I_110 and TAF1D/TAF_I_41, serves as the foundation for the creation of a proficient initiator complex on the promoter (Learned et al., 1985)(Eberhard et al., 1993)(Gorski et al., 2007)(Comai et al., 1994). RRN3, in turn, facilitates the recruitment of RNA Pol I to the promoter and serves as a bridge between RNA Pol I and the SL1 complex anchored at the promoter. Therefore, the SL1-RNA Pol I-RRN3 ternary complex anchors at the promoter bound by UBF, leading to the activation of rDNA transcription (Friedrich et al., 2005).

RNA Pol I mediates its transcriptional activity at the rDNA locus by repeatedly engaging on tandem repeats of rDNA. Significantly, not all repeats exhibit transcriptional activity; nearly half of them are maintained in a transcriptionally poised state, primarily through epigenetic mechanisms like histone (H3K9me, H4K20me) or DNA methylations (McStay & Grummt, 2008). The rDNA that is actively transcribed is typically linked with euchromatin, exhibiting hypomethylation of rDNA and carrying histone modifications commonly associated with gene activation, such as H3K4 methylation and H3K9 acetylation (McStay & Grummt, 2008). Examination of rDNA chromatin structure through genomic analysis has unveiled the existence of H3K4 methylation marks at both rDNA promoters and intergenic regions (Zentner et al., 2011). Nevertheless, the specific role played by H3K4 methylation in the activation of rDNA transcription remains unclear. The existence of H3K4 methylation marks specifically at both promoters and intergenic regions of rDNA repeats suggests that the H3K4 lysine methyltransferase (KMT) complex could have a significant role in regulating rDNA transcription. But how this regulation is brought about by the H3K4 KMTs is not known.

H3K4 methylation is mediated by the COMPASS/Mixed Lineage Leukemia (MLL)/lysine methyl transferase 2 (KMT2) family of proteins. The mammalian COMPASS family consists of six members— MLL1/KMT2A, MLL2/KMT2B, MLL3/KMT2C, MLL4/KMT2D, SET1A/KMT2F and, SET1B/KMT2G — each actively engaged in complexes that include four common subunits collectively referred to as WRAD (WDR5, RBBP5, ASH2L, and DPY30), (Shilatifard, 2012)(Ernst & Vakoc, 2012) (Sugeedha et al., 2021). These family members modulate transcription primarily via their Su(var)3-9, Enhancer-of-zeste, Trithorax (SET) domain, though some members like KMT2A and KMT2B also possess transcriptional activation domain (TAD). Interestingly, KMT2 family members exhibit variations in their methyl transferase activity. KMT2F is the global KMT, mainly responsible for depositing H3K4me3 marks on promoters throughout the genome. KMT2A and KMT2B, on the other hand, demonstrate selective H3K4me3 activity at specific loci, such as Hox loci (Li et al., 2022)(Wu et al., 2008)(Denissov et al., 2014). Several reports suggest that H3K4 methylation deposited by the members of the COMPASS family plays an important role in facilitating the assembly of the transcription pre-initiation complex and the recruitment of RNA polymerase II (RNA Pol II) to specific gene promoters (Wang et a l., 2009)(Laub erth et a l., 2013). Recent study depleted shared subunits within the COMPASS family to show that the abrupt decrease in H3K4me3 shows no discernible effects on the transcriptional initiation of RNA Pol II; nevertheless, it leads to a widespread decrease in transcriptional output, an increase in RNA Pol II pausing, and a slowdown in elongation, underscoring the crucial role of H3K4 methylation, mediated by the KMT2 family, in modulating RNA Pol II activity (H. Wang et al., 2023)(Hu et al., 2023). While there is increasing evidence supporting the role of H3K4 methylation in regulating RNA Pol II activity, the role of H3K4 methylation in connection with RNA Pol I regulation remains largely unknown.

Here, we undertook studies to understand how RNA Pol I transcription is regulated by the H3K4 KMT enzymes. We made use of customized genome assemblies for rDNA to demonstrate that KMT2A and KMT2F bind to entire rDNA loci (George et al., 2023). The binding of these enzymes correlated closely with the H3K4me2 and H3K4me3 marks. In order to demonstrate direct association, we used biochemical assays to examine the interaction between common subunit, WDR5, as well as KMT2A and KMT2F with the members of RNA Pol I machinery. Our transcriptional assays revealed reduction in rRNA levels upon depletion of KMT2 members. Through mutational analysis, we discovered that KMT2F functions through its SET domain to regulate rDNA transcription. Our knock down studies substantiated that KMT2F is the primary enzyme responsible for regulating the deposition of H3K4me3 on rDNA. Subsequently, we investigated the impact of perturbing the epigenetic landscape of rDNA, and the formation of the RNA Pol I PIC, using ChIP assays. Our findings indicate that methyltransferase activity of KMT2F is requisite for PIC formation, and ensuing transcription by RNA Pol I. Thus, our results reveal H3K4me3 mark as a key determinant in RNA Pol I PIC formation and active transcription from this locus.

## Results

### KMT2A and KMT2F bind to the ribosomal DNA (rDNA) locus

Nucleolus is a distinct structure and site of transcription and processing of rRNA. Proteins involved in rRNA transcription are often enriched at the nucleolus. We and others have previously reported that KMT2A localized to the nucleolus in human cells (Karole et al., 2018)(Yano et al., 1997)(Caslini et al., 2000). We undertook further experiments here to interrogate if KMT2A has a functional role in rRNA transcription. As KMT2F is the global H3K4 KMT, we included it in our analysis here. Using immunofluorescence staining (IFS) we observed that KMT2A and KMT2F co-localized with nucleolar marker protein B23 in U-2OS cells (Figure 1A). Specific siRNAs directed against KMT2A or KMT2F transcripts reduced nucleolar localization of both proteins confirming specificity of this localization (Supplemental Figure S1A-B). We further performed IFS on various cell lines including MCF-7, HeLa and non-transformed IMR90-tert, and observed that both proteins localized to the nucleolus in these cells (Supplemental Figure S1 C-D).

**Figure 1:**
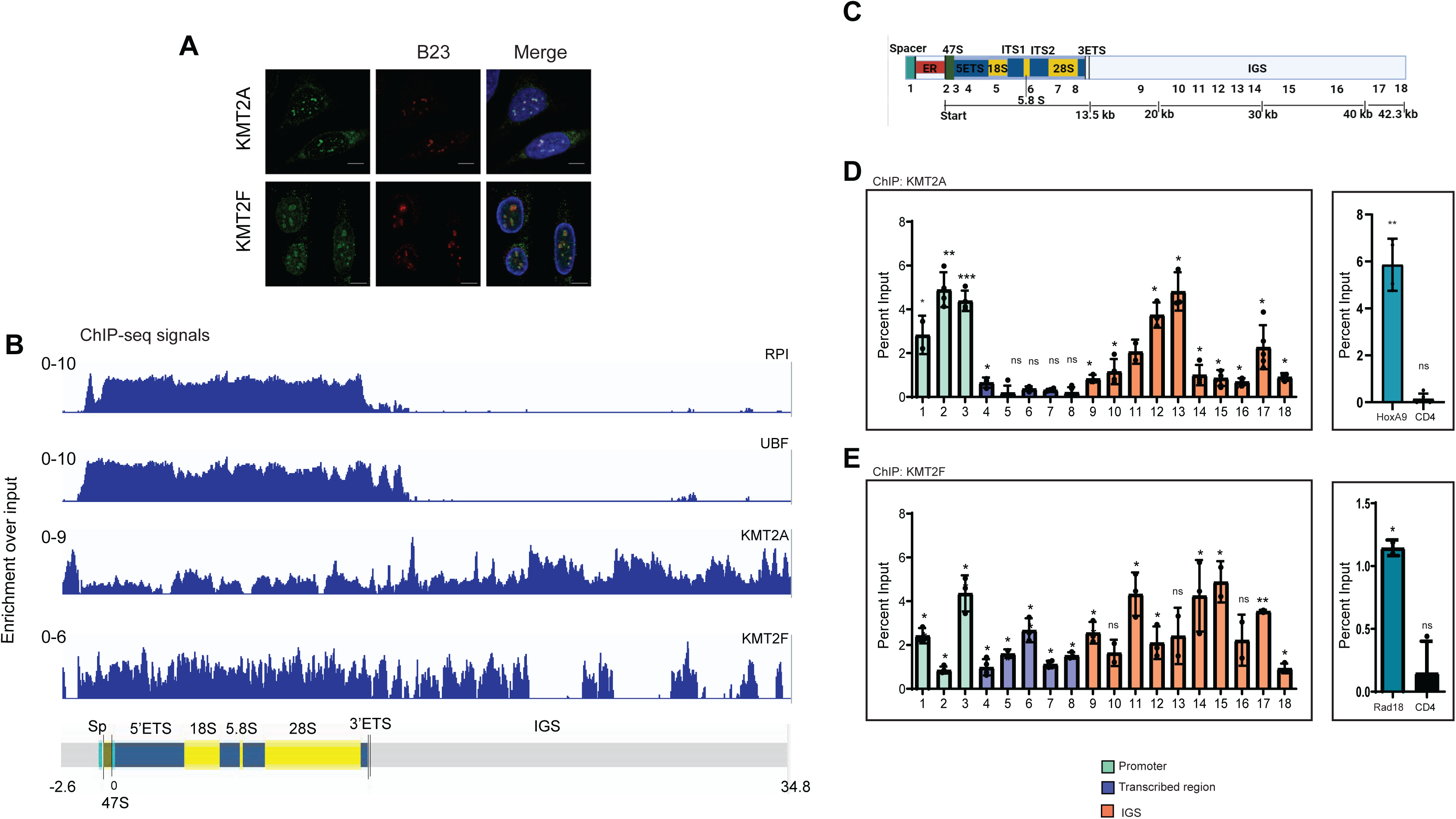
KMT2A and KMT2F bind to the human ribosomal DNA (rDNA) repeats. **A.** Immunofluorescence staining (IFS) of endogenous KMT2A and KMT2F is shown.U-2OS cells were co-stained with either KMT2A or KMT2F and nucleolar marker B23. DNA was counterstained with 4,6-diamidino-2-phenylindole (DAPI, blue). Scale bar, 10 μm. **B.** Chromatin immunoprecipitation-sequencing (ChIP-seq) maps, illustrating the binding of RNA Pol I (RPA194), UBF and KMT2’s at the human rDNA locus, are shown. ChIP enrichment for each factor is expressed as the ratio of immunoprecipitated ChIP-seq signal to Input DNA ChIP-seq signal. The vertical scale (log2) in each panel indicates the enrichment of each component relative to the Input DNA dataset. On the X axis, tracks were truncated at 2.6 kb upstream of the main promoter (47S), denoted as −2.6 (kb), and at 34.8 (kb) downstream (also see Methods and Supplemental Figure S2). **C.** Schematics representing 43kb human rDNA repeat drawn to scale. Each human rDNA repeat unit contains a 13kb sequence encoding the precursor ribosomal RNA, and Intergenic spacer region (IGS) spanning 30 kb. Primers (1 to 18) used for qRT-PCR represent different regions of the rDNA repeat. Co-ordinates and sequence of each primer is given in supplementary Table S1. Spacer: spacer promoter; ER: Enhancer repeat; 47S: core promoter; ETS: External transcribed spacer; ITS: Internal transcribed spacer. **D-E.** ChIP analysis of KMT2A (**C**) and KMT2F(**D**) is shown. Primers 1-18 were used to assess the binding of KMT2A/KMT2F over the 43kb human rDNA repeat. Spacer promoter, terminator element (T0) and 47S (core) promoter are represented by primers 1-3 respectively; transcribed region: 4-8 and IGS by 9-18. *HOXA9* and *RAD18* were used as positive control regions for KMT2A and KMT2F binding respectively. *CD4* was used as negative control for KMT2A and KMT2F ChIP. Data is presented from three or more experiments. Error bars represent SD. \**P* ≤ 0.05, ** *P* ≤ 0.005, \*\*\**P*≤ 0.0005, \*\*\*\**P*≤ 0.00005, ns: not significant *P* > 0.05 (two-tailed Students *t* test). Green: RNA pol I promoter; blue: RNA pol I transcribed region; orange: IGS.

As rDNA are repetitive in nature, they have been left unannotated in most genome assemblies. However, the rDNA repeat is a large locus to be mapped with chromatin immunoprecipitation (ChIP) alone. In order to interrogate whether the KMT2 proteins bind to rDNA repeats or not, we performed paired-end chromatin immunoprecipitation followed by sequencing (ChIP-seq) for KMT2A and KMT2F. The reads obtained were mapped to custom human genome assembly in which a single unit of rDNA has been added (Figure 1B-C) (George et al., 2023). We included the ChIP-seq analyses of RNA Pol I largest subunit, RPA 194 (referred to as RNA Pol I in the Figure 1B) and UBF as positive control (Figure 1B). Both RNA Pol I-and UBF-bound reads mapped to the transcribing unit of rDNA as reported previously (Figure 1B, Supplementary Figure S2; (Zentner et al., 2011)(Herdman et al., 2017). We observed that KMT2A and KMT2F exhibited binding to the entire region of rDNA sequence including the RNA Pol I promoter, transcribed unit and the IGS (Figure 1B, Supplementary Figure S2). KMT2F showed a consistent binding in the RNA Pol I transcribed unit, with distinctive peaks in the IGS whereas KMT2A bound throughout the RNA Pol I transcribed region but displayed higher peaks with continuous binding in the IGS region. To strengthen our confidence in the binding of these proteins to the rDNA locus, we utilized data from other KMT2 family-associated proteins. The WRAD complex plays a critical role in determining the binding specificity of KMT2 family members on chromatin (Bochyńska et al., 2018). For this we took the advantage of ChIP-seq data for WDR5, a common component shared by all KMT2 family members (Thomas et al., 2015). Upon analyzing the ChIP-seq tracks, we observed significant WDR5 binding in the transcribed unit of the rDNA, along with a distinct peak in the IGS region (Supplementary Figure S2). Since KMT2F lacks direct DNA-binding domains, it is recruited to chromatin via the WDR82 protein (Lee & Skalnik, 2008). WDR82, which uniquely binds to the KMT2F complex, was further analyzed using ChIP-seq data (Park et al., 2022) to explore its binding pattern across the entire rDNA locus. We found that WDR82 showed higher enrichment in the transcribed unit (Supplementary Figure S2), consistent with KMT2F binding. Thus, our ChIP-seq data reveals subtle differences in binding of KMT2A and KMT2F at the rDNA locus.

The above results gave us confidence that the KMT2 complexes bind to rDNA. However, as rDNA genome assemblies are still evolving, we validated our findings by performing ChIP analyses using select primers spanning the entire rDNA unit (Figure 1C, Supplementary Table 1). Primer #1 corresponded to spacer promoter, #3 to the core 47S promoter, # 4-7 to gene body, #8 to termination repeats and #9-18 spanned the IGS as shown (Figure 1C, Supplementary Table 1). We also used canonical targets as positive control (HoxA9 for KMT2A and Rad18 for KMT2F) and CD4 as negative control. We first performed ChIP with RNA Pol I and UBF (Supplementary Figure S3A-B). As expected, both proteins showed highly enriched binding on the rDNA promoter and transcribed region. This binding decreased abruptly after the termination signal (primer #8; Supplementary Figure S3A-B). The results obtained here gave us confidence in our protocol and indicated that we are able to ChIP for proteins bound to rDNA successfully. We observed that in directed ChIP, KMT2A was significantly enriched at the RNA Pol I promoter and IGS (Figure 1D), whereas KMT2F bound primarily in the region of RNA Pol I promoter, and transcribed unit (Figure 1E). In order to asses if the role of these proteins is cell-type specific, we performed ChIP in non-transformed IMR90-tert cells, fibroblasts isolated from normal lung tissue and immortalized using human Telomerase Reverse Transcriptase (hTERT). Similar to our observations made in HEK293 cells (Figure 1D-E), KMT2A displayed binding primarily at the RNA Pol I promoter and IGS, whereas KMT2F bound throughout the rDNA repeat (Supplementary Figure S3C-D). Taken together, our data indicates that KMT2A and KMT2F bind to rDNA repeat locus including the RNA Pol I promoter.

### rDNA locus is enriched for H3K4me2 and H3K4me3 mark

To understand if the binding of KMT2 proteins has a functional role on the rDNA locus, we performed similar analyses on previously published paired end ChIP-seq data sets for H3K4me2 and H3K4me3(Liu et al. 2022). We observed that H3K4me2 and H3K4me3 marks were present on the RNA Pol I promoter and transcribed region. This binding continued in the IGS region eventually getting confined to distinct peaks (Figure 2A). When aligned together, H3K4me2 and H3K4me3 peaks seemed to correlate more to KMT2F binding (Figure 2A, S2). Again, we validated our findings by ChIP analyses and observed consistently high enrichment of these marks on RNA Pol I promoter and IGS region in HEK293 (Figure 2B-C). Consistent with the presence of these marks, H3 could be detected in the entire rDNA repeat region (Supplemental Figure S3E).

**Figure 2:**
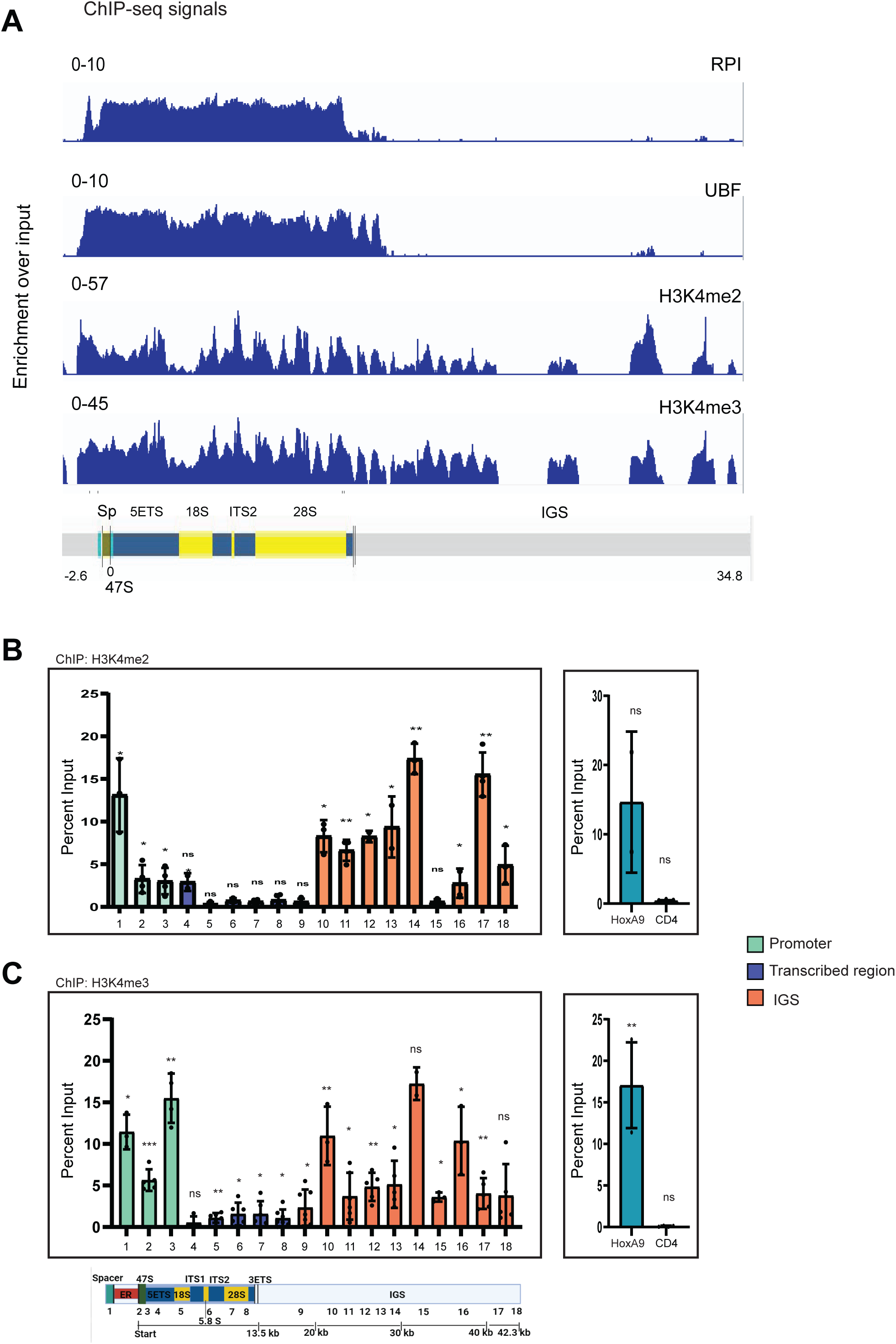
Active Epigenetic signature of human rDNA. **A.** ChIP-seq maps illustrating the binding of RPA194, UBF and distribution of active epigenetic marks (H3K4me2 and H3K4me3) at the human rDNA locus are shown. ChIP enrichment for each factor is analysed as in Figure 1B. *Please note that RPA194 and UBF tracks are duplicated from* Figure 1B *and shown here for ease of comparison.* (also see Supplemental Figure S2) **B,C**. Enrichment of H3K4me2 (**B**) and H3K4me3(**C**) across human rDNA as revealed by ChIP followed by qRT-PCR. Primers labelled from 1-18 were used to access distribution pattern of H3K4me2 and H3K4me3 across 43kb repeat. *HOXA9* and *CD4* were used as positive and negative control regions respectively. Data shown as means ± SD from three or more than three individual biological replicates. Error bars represent SD. \**P* ≤ 0.05, ** *P* ≤ 0.005, \*\*\**P*≤ 0.0005, \*\*\*\**P*≤ 0.00005, ns: not significant *P* > 0.05 (two-tailed Students *t* test). Green: RNA pol I promoter; blue: RNA pol I transcribed region; orange: IGS

### H3K4 KMT complex interacts with RNA Pol I machinery

In order to establish a regulatory role of our KMTs in RNA Pol I mediated transcription, we tested if these complex specifically interacts with RNA Pol I machinery. RNA Pol I holo-complex is composed of 14 subunits, out of which 7 are unique to RNA Pol I (hereafter called RNA Pol I specific subunits; Figure 3A). These include RPA 194, RPA 116, RPA12, RPA49, RPA34, RPA43 and RPA14 (RPA 14 was discovered in yeast however it is not yet been characterized in mammals; (Viktorovskaya & Schneider, 2015). The remaining subunits are shared between all RNA polymerases. We tried to express all six RNA Pol I specific subunits as N-terminal Glutathione S-transferase (GST) fusions in bacteria. However, technical challenges limited us to successful expression of RPA12, RPA43, RPA49, and RPA34. Interestingly, three of these subunits are classified as the peripheral subunits of RNA Pol I and have been implicated in RNA Pol I initiation and elongation ( Kuhn et al., 2007). We tested *in vitro* interactions between bacterially expressed GST-tagged RNA Pol I specific subunits with bacterially expressed WDR5 (Figure 3B). We used GST-KMT2A SET domain as positive control and GST alone as negative control. We observed that WDR5 specifically interacted with RPA49 and RPA34 (Figure 3B). As RPA49 and RPA34 showed consistent interaction with WDR5, we choose these subunits to pull down the SET-domain containing subunits of H3K4 KMT enzymes. Remarkably, GST-RPA49 and GST-RPA34 could pull-down the high molecular weight proteins KMT2A and KMT2F robustly, indicating that our proteins specifically interacted with the RNA Pol I subunits in the cells (Figure 3C).

**Figure 3:**
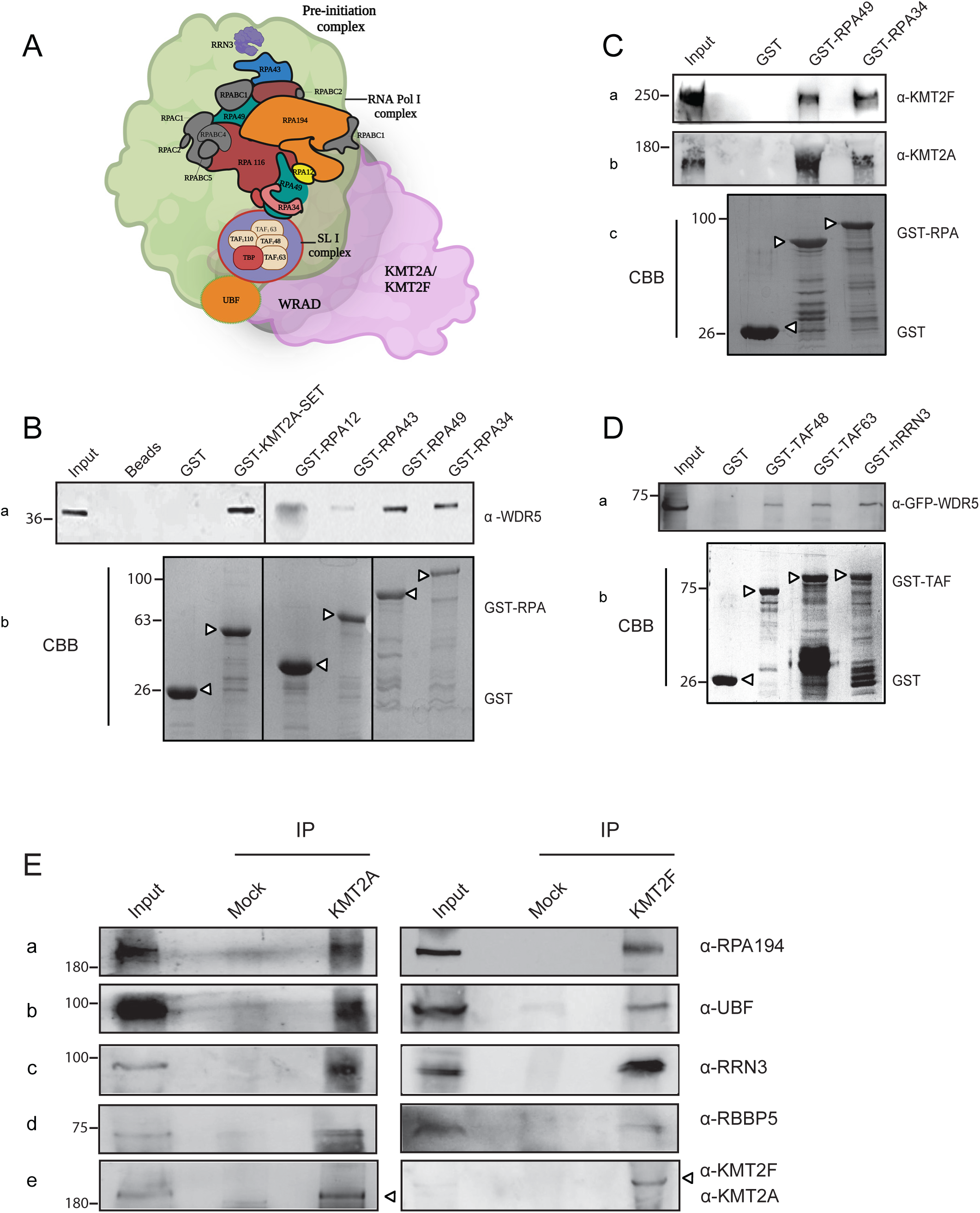
RNA Pol I machinery interacts with KMTs. **A.** Schematic showing the arrangement of RNA Pol I holo complex with its associated proteins. RNA Pol I has 14 subunits, out of which 7 are shared with Pol II and Pol III while rest are unique to Pol I. Unique subunits, shown in colour. RPA190, RPA116 and RPA12 forms the catalytic core of the RNA Pol I whereas RPA34, RPA43 and RPA49 are classified as the peripheral subunits. UBF prepares the rDNA promoter for RNA Pol I docking. This is followed by binding of the SL1 complex, which consists of TATA binding protein (TBP) and four transcription-associated factors (TAFs) TAF1A/TAFI48, TAF1B/TAFI63, TAF1C/TAFI110 and TAF1D/TAFI41. hRRN3 identifies and recruits free RNA Pol I to the rRNA gene promoter that is already bound by UBF and SL1 complex. Based on our results below, we have put forward a model, where KMT2A/2F along with WRAD complex, interact with RNA Pol I complex. **B.** Bacterially expressed GST, GST-KMT2A-SET domain, GST-RPA12, GST-RPA43, GST-RPA49, and GST-RPA34 were incubated with the bacterially expressed and purified WDR5. The interaction was visualized by Western blot by using anti-WDR5 antibody. *Please note that WDR5 and GST-RPA12 bands co-migrate and we believe that the smudge seen in GST-RPA-12 lane is produced by large amount of GST-RPA-12 and not WDR5*. **C.** Bacterially expressed GST and GST-tagged RPA49 and RPA34 were used to assess the interaction of KMT2A and KMT2F with RNA Pol I complex. This interaction was confirmed by immunoblotting with α-KMT2F and α-KMT2A antibodies. **D.** GST, GST-RRN3 and GST tagged SL1 complex proteins (TAF48 and TAF63) were bacterially purified and incubated with whole cell extracts from cells ectopically expressing GFP-WDR5 (endogenous WDR5 could not be probed due to technical difficulties). The interaction was confirmed by immunobloting with α-GFP. (**B**-**D**)The amount of protein taken for GST pull down assay, stained with Coomassie Brilliant Blue (CBB), is shown at the bottom. Relevant protein bands are indicated with arrow heads. **E.** Nuclear extract prepared from HeLa cells was subjected to endogenous immunoprecipitation (IP) using antibodies against KMT2A and KMT2F. The anti-immunoglobulin G (IgG)-antibody was used as control (Mock). The immunoblot was probed with α-RPA194, α-UBF, α-RRN3 α-RBBP5, and α-KMT2A or α-KMT2F antibodies. RBBP5, was used as a positive control for anti-KMT2A and anti-KMT2F IP. (**B**-**E**) The molecular weight marker is indicated on the left.

In addition to the RNA Pol I enzyme, rDNA transcription requires UBF, SL1 complex and transcription initiation factor hRRN3 (McStay & Grummt, 2008). SLl-RNA Pol I-hRRN3 ternary complex docks at UBF bound RNA Pol I promoter that results in activation of transcription (Gorski et al., 2007). In order to establish that the KMTs interact with transcription-competent RNA Pol I complex, we checked for interaction of TAF 48 and TAF 63 (subunits of SLI complex) and hRRN3, with WDR5. Consistent with our hypothesis, WDR5 interacted with SL1 complex and hRRN3 (Figure 3D). We could further prove that KMT2A and KMT2F interacted with various components of RNA Pol I by endogenous immunoprecipitation experiments where we pulled down KMT2A or KMT2F proteins using specific antibodies and verified their interaction with RPA194, UBF and hRRN3 (Figure 3E, S4A). Altogether, our results show that KMT2A and KMT2F interact with RNA Pol I and multiple components its transcription machinery.

### KMT2F regulates rRNA transcription via its SET domain

We have shown so far that the H3K4 KMTs— KMT2A and KMT2F— bind to rDNA locus and interact with RNA Pol I transcription machinery. Given these results, we hypothesized that these KMTs activate transcription of rRNA. rRNA is transcribed into 13 kb long 47S precursor ribosomal RNA (pre-rRNA) which gets processed at its 5’ end, called 5’ external transcribed spacer (5’ETS) as soon as it gets transcribed to give rise to 45S pre-RNA (McStay & Grummt, 2008). Therefore, the 5’ETS can be used to detect the transcript level of pre-rRNA by quantitative real-time PCR (qRT-PCR). We used RNAi to independently knock down KMT2A, KMT2F and WDR5 in U-2OS cells. The transcript levels of KMT2A, KMT2F and WDR5 decreased by more than 60% and we observed a corresponding decrease in the 5’ETS levels in these samples (Figure 4A, S4 B-D) indicating that KMT2A and KMT2F are required for rDNA transcription.

**Figure 4:**
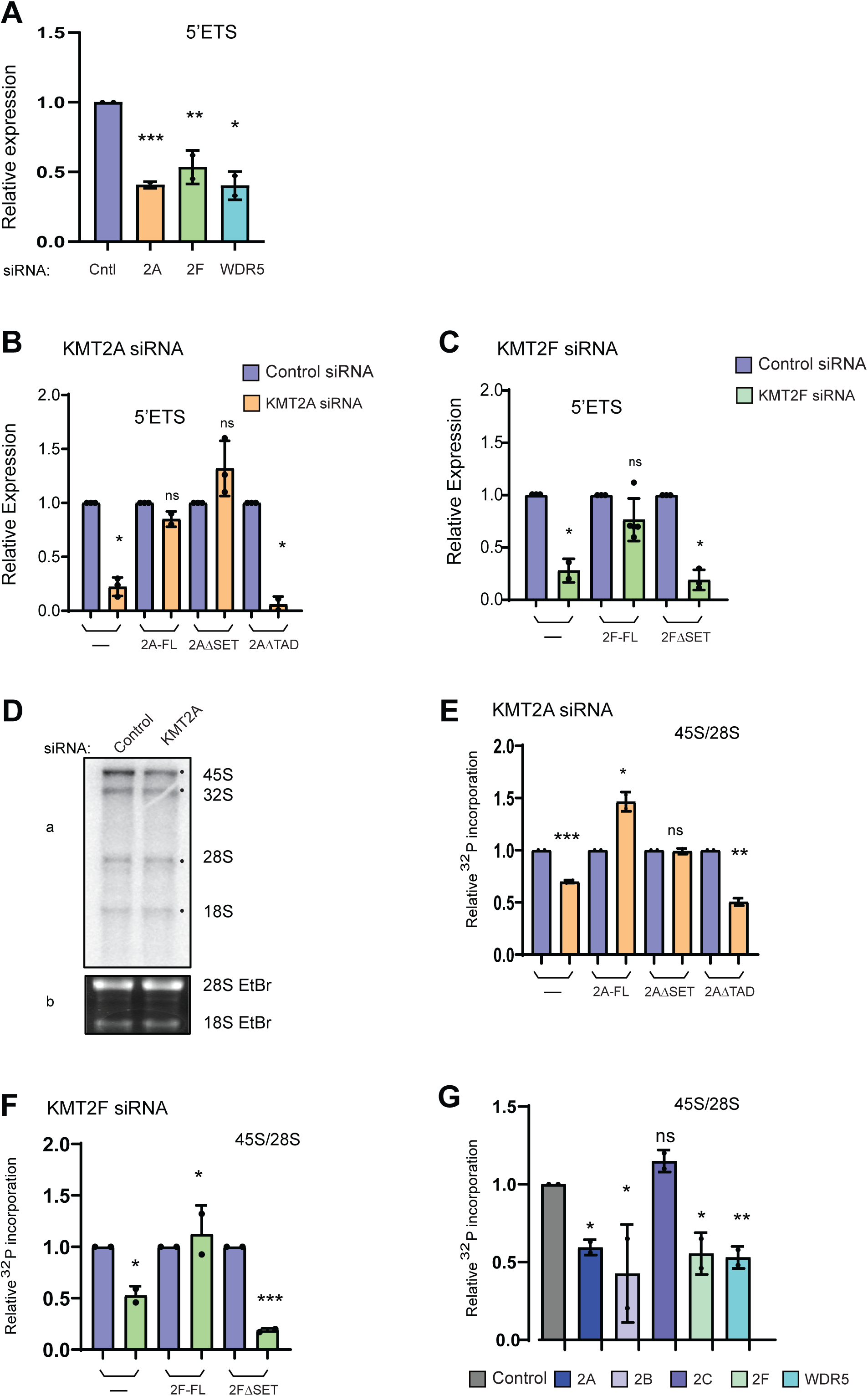
RNAi mediated depletion of MLLs affect ribosomal DNA transcription. **A.** qRT-PCR analysis of 5 ETS transcript levels after treating the cells with Control, KMT2A, KMT2F and WDR5 siRNA is shown. Error bars denote SD. *P ≤ 0.05, ** P ≤ 0.005, ***P≤ 0.0005, ns: not significant P > 0.05 (two-tailed Students *t* test). **B,C.** qRT-PCR analysis of 5’ ETS transcript levels in various KMT2A and KMT2F mutant cell lines treated with specific siRNAs is shown. **B**. KMT2A depleted cells were complemented with stably expressing KMT2A full length (2A-FL), KMT2AΔSET (2AΔSET), and KMT2AΔTAD (2AΔTAD) cells and5’ ETS transcript levels were analysed as shown. **C.** Shows 5’ ETS transcript levels in control cells, KMT2F full length (2F-FL), and KMT2FΔSET (2FΔSET) cells. Error bars denote SD. *P ≤ 0.05, ** P ≤ 0.005, ***P≤ 0.0005, ns: not significant P > 0.05 (two-tailed Students *t* test). **D.** Representative autoradiogram showing the ^32^P orthophosphate labelled ribosomal RNA upon loss of KMT2A. U-2OS cells were treated with Control and KMT2A siRNA. Extracted total RNA was run on agarose gel and visualized by autoradiography (panel a). Ethidium bromide stained 28S RNA levels were determined by running a formaldehyde agarose gel (panel b). **E.** Effect of mutational analysis of KMT2A on rRNA transcription. Densitometry quantification of 32P incorporated 45S rRNA levels in endogenous KMT2A depleted Control (U-2OS), KMT2A full length (2A-FL), KMT2AΔSET (2AΔSET) and KMT2AΔTAD (2AΔTAD) cells is shown. The relative intensity was measured by normalizing 45S levels with 28S rRNA loading control. Each value is an outcome of two independent experiments. S.D is represented by error bars. The statistical significance was calculated by two-tailed student *t* test *P ≤ 0.05, **P ≤ 0.005, ns-not significant P > 0.05. **F.** Effect of mutational analysis of KMT2F on rRNA transcription. Shown is densitometry quantification of ^32^P incorporation into 45S rRNA levels in endogenous KMT2F depleted U-2OS cells which were complemented with stably expressing KMT2F full length (2F-FL), KMT2FΔSET (2FΔSET). **G.** The 45S rRNA transcript levels upon loss of different KMT2 family members is shown. U-2OS cells were treated with KMT2A, KMT2B, KMT2C, KMT2F and WDR5 siRNAs. (**E**-**G**) ^32^P incorporation into 45S rRNA levels were measured by densitometry. The relative intensity was measured by normalizing 45S levels with 28S rRNA loading control. Each value is an outcome of two independent experiments. S.D is represented by error bars. The statistical significance was calculated by two-tailed student *t* test *P≤ 0.05, **P ≤ 0.005, ns-not significant P > 0.05.

To investigate the specificity of the observed reduction of 5’ETS in KMT2A and KMT2F siRNA, we conducted complementation assays where we examined the rRNA transcripts in siRNA-treated cells that were expressing respective recombinant full-length protein. Additionally, we utilized SET-domain deleted KMT2A/F protein(s) to gain insights into the role of the SET domain in rRNA transcription (Supplemental Figure S4E-F). For this purpose, we used stable cell lines created in U-2OS cells (Ali et al., 2014)(Malik et al., 2023). In order to specifically target the endogenous KMT2A or KMT2F transcript with our siRNA treatment, and avoid affecting the recombinant transcript, we (i) designed siRNA to target the 3’ untranslated region (UTR) of the precursor mRNA for KMT2A; (ii) created recombinant siRNA-resistant KMT2F constructs by introducing silent mutations (as shown in Supplemental Figure S4F).

We observed a substantial reduction in KMT2A and KMT2F transcript levels following treatment of U-2OS cells with KMT2A or KMT2F siRNAs, which was largely rescued in their respective stable cell lines (Figure 4B-C, S4G-H). Corresponding to the reduction in KMT2A levels in U-2OS cells, as before, we observed a significant decrease in 5’ETS levels in the siRNA-treated U-2OS cells. This reduction was rescued in cells overexpressing full-length KMT2A and KMT2AΔSET but not in KMT2AΔTAD cells (Figure 4B, S4G), indicating that KMT2A regulates rDNA transcription through its TA domain. On the other hand, 5’ETS transcript levels were rescued in cells expressing full-length KMT2F, but not in cells expressing KMT2FΔSET, suggesting that KMT2F regulates rDNA transcription via the methyltransferase activity of its SET domain (Figure 4C, S4H).

In order to validate our results from above experiments, we used an assay to measure ongoing rRNA transcription where we gave a short pulse with [32P] orthophosphoric acid to label cells after RNAi treatment, extracted and ran RNA on agarose gel and visualized 47/45S pre-rRNA by autoradiography (Nguyen & Mitchell, 2013). The assay enabled us to visualize freshly processed 32S, 28S and 18S rRNA moieties though 45S rRNA was the predominant species seen (Figure 4D). We also used ethidium bromide gel to visualize 28S RNA as loading control, as it is largely made during previous rounds of transcription and remains stable over three days. Our results from pulse-chase studies concurred with our observations made with qRT-PCR (from Figure 4A-C) and we observed a consistent decrease in the 45S rRNA upon knock down of KMT2A (Figure 4D-E,G, S5A,C) and KMT2F (Figure 4F-G, S5B,C). This decrease was reproduced in HeLa, MCF-7 and IMR90-tert cell lines indicating that KMT2A and KMT2F regulate 45S RNA irrespective of cell type (Supplemental Figure S5D-E). We observed that while both KMT2A and KMT2F play a specific role in regulating rRNA transcription, only KMT2F did so in a SET-domain dependent manner (Figure 4E-F, S5A-B). In contrast, loss of TA but not SET domain in KMT2A resulted in decreased rRNA transcription (Figure 4E, S5A). Taken together our results suggest that KMT2A and KMT2F use divergent mechanisms to regulate rRNA transcription.

### Select members of KMT2 family regulate rRNA transcription

Our results above prompted us to check if other members of KMT2 family had a role in transcriptional regulation of rRNA. We depleted various KMT2 members by RNAi and checked for relative levels of 45S rRNA. Our results show that besides loss of KMT2A and KMT2F, loss of KMT2B but not KMT2C, resulted in decrease in 45S rRNA levels (Figure 4G, S5C). Consistent with these findings, we could detect KMT2B in the nucleolus (Supplemental Figure S5F). Our results suggest that only select members of KMT2 family engage with RNA Pol I to regulate rRNA transcription.

### KMT2F is the H3K4me3 methyltransferase for rDNA locus

Several reports indicate that rDNA bears the H3K4me3 marks and these marks are involved in shaping the epigenetic landscape of this loci (Zentner et al., 2011)(W. Wang et al., 2011)(Feng et al., 2010). Our data indicates that some members of the KMT2 family regulate rRNA transcription, and we have shown that KMT2F utilizes its SET domain to regulate rRNA transcripts. Further, it binds on the rDNA and interacts with RNA Pol I. To investigate the role of this protein in detail, we used shRNA mediated knock down for KMT2F as described previously (Malik et al., 2023). Consistent with the reduced protein levels on the blots, the chromatin binding of KMT2F was drastically reduced on rDNA transcribed by RNA Pol I (Figure 5A-B, S6A). Remarkably, consistent with the role of SET domain of KMT2F in our rRNA transcript analyses, knock down of KMT2F resulted in drastic reduction of H3K4me2 and H3K4me3 levels on the whole rDNA loci (Figure 5C-D, S6B-C). Although we reported statistically significant levels of H3K4me2 only on the promoter region of rDNA (Figure 2C), the loss of KMT2F resulted in the reduction of H3K4me2 levels throughout RNA Pol I transcribed region when compared with control (Figure 5C). Our results indicate that KMT2F is the primary H3K4 methyltransferase of rDNA.

**Figure 5:**
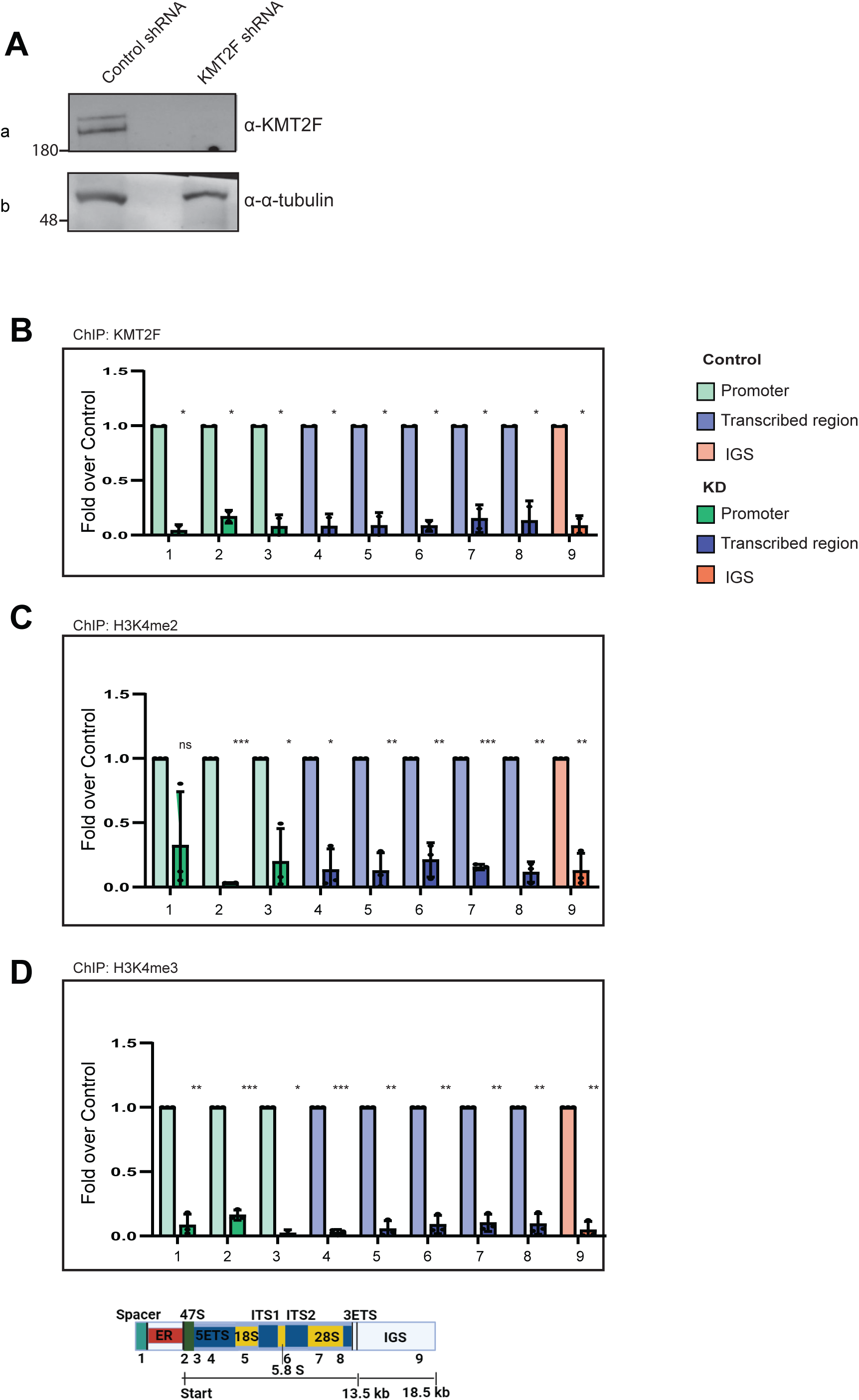
KMT2F deposits H3K4me3 marks to regulate epigenetic state of human rDNA. **A** Immunoblots of whole cell extracts were performed on scrambled shRNA (control shRNA) and KMT2F shRNA treated cells. Immunoblots were probed with α-KMT2F and α-α-tubulin as shown. **B-F**. ChIP qPCR of KMT2F (**B**) H3K4me2 (**C**) H3K4me3 (**D**) H3K9ac (**E**) H3K9me3 (**F**) in control shRNA and KMT2F shRNA conditions. Data is presented as fold over control. Experiments were performed two (for KMT2F) or more than two times as biological replicates. Error bars represent SD. \**P* ≤ 0.05, ** *P* ≤ 0.005, \*\*\**P*≤ 0.0005, \*\*\*\**P*≤ 0.00005, ns: not significant *P* > 0.05 (two-tailed Students *t* test).

### Loss of KMT2F HMTs affects epigenetic landscape of rDNA

H3K4me2 and H3K4me3 marks are associated with positive transcription and accordingly influence the surrounding chromatin epigenetic landscape (Liu et al., 2016)(Sankar et al., 2020). H3K9 acetylation (H3K9ac) is another mark associated with open chromatin. Therefore, we tested the levels of H3K9ac on rDNA in KMT2F KD. Loss of KMT2F resulted in reduced levels of H3K9ac (Figure 5E, S7A). Consistent with the loss of active marks like H3K9ac and H3K4me3, loss of KMT2F was accompanied by 5-10-fold increase in H3K9me3 mark (Figure 5F, S7B), indicating that by depositing H3K4me3 mark, KMT2F prevents the heterochromatinization of rDNA and keeps it open for transcription.

### Loss of KMT2F abrogates pre-initiation complex formation by RNA Pol I

Loss of KMT2F decreased rRNA transcription and impacted the rDNA epigenetic landscape adversely. In order to find out exactly what step of RNA Pol I transcription is affected; we undertook further studies. The human rDNA transcription involves following sequence: rDNA is activated when UBF binds to the RNA Pol I promoter. Then UBF recruits the SL1 complex to form the PIC at the rDNA promoter. Following this, UBF interacts with RNA Pol I through its RPA34 and RPA49 subunits, while SL1 interacts with the transcription factor RRN3. All factors bind to 47S promoter as well as spacer promoter. Once the PIC formation stabilizes RNA Pol I, RRN3 is released from the RNA Pol I complex, and RNA Pol I proceed with transcript elongation to yield 47S pre-rRNA.

In KMT2F KD, we observed that the occupancy of RNA Pol I was drastically reduced all across the transcribed region (Figure 6A). Similarly, the binding of both SL-1 complex and RRN3 to rDNA promoter was highly reduced when compared to control samples (Figure 6B-C). We observed a 50% decrease in UBF binding on KMT2F KD on the spacer promoter and 47S promoter, which reduced further in the body of the transcribed unit (Figure 6D). We have shown that KMT2F deposits H3K4me3 marks to keep the rDNA open for transcription (Figure 5). Further, we observe that RNA Pol I pre-initiation complex formation is completely abrogated upon loss of KMT2F resulting in reduced rRNA transcription. Taken together, our data suggests that H3K4me3 marks deposited by KMT2F is requisite for successful PIC formation and transcription by RNA Pol I at the rDNA locus.

**Figure 6:**
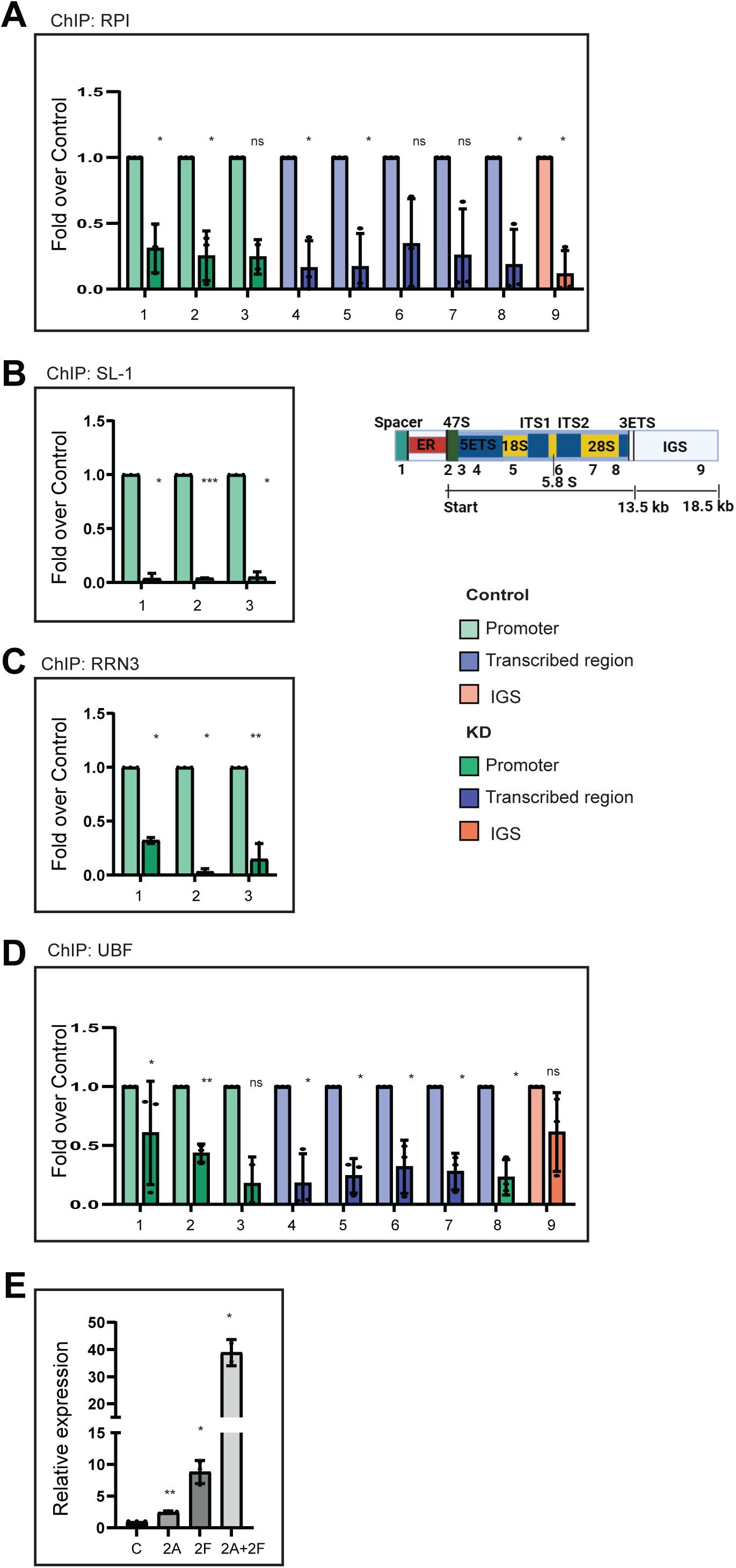
Loss of KMT2F affect the formation of RNA Pol I pre-initiation complex on rDNA. **A-D** ChIP followed by qRT-PCR of RNA Pol I (**A**), SL1 represented by TAF1C (**B**), RRN3 (**C**), and UBF (**D**) was performed in cells transfected either with scrambled or KMT2F shRNA. Fold over control was determined by dividing the ChIP signals of KD cells by ChIP values of control cells for indicated antibodies. Promoter (#1-3; green), transcribed region (#4-8, blue) and IGS (9; orange) primers were used to quantify ChIP qPCR signals. Error bars represent mean ± SD of three or more than three individual biological replicates. \**P* ≤ 0.05, ** *P* ≤ 0.005, \*\*\**P*≤ 0.0005, \*\*\*\**P*≤ 0.00005, ns: not significant *P* > 0.05 (two-tailed Students *t* test). **E.** KMT2A (2A), KMT2F(2F) were overexpressed alone or together (2A+2F) along with vector control (C) in HEK293 cells. Total RNA was prepared from over expressed cells followed by cDNA synthesis. 5 ETS transcript levels were quantified in all the three cases using primer #3. n=3 \**P* ≤ 0.05, ** *P* ≤ 0.005 not significant *P* > 0.05 (two-tailed Students *t* test).

### Recruitment of the KMT2F to the rDNA locus

KMT2F lacks the direct DNA binding domains. However, it is indirectly recruited to the chromatin through its association with various proteins. CFP1 acts as the primary targeting component for the KMT2F complex, recruiting it to CpG island promoters, while WDR82, alone or in association with other factors, targets KMT2F to transcription start sites (Brown et al. 2017) (Lee & Skalnik, 2008); (Franks et al. 2017). Other recruitment pathways have been described for specific context like cell cycle-regulation (Tyagi et al., 2007). Hence, KMT2F is engaged in multivalent interactions with the chromatin. As the presence of KMT2F on the RNA Pol I promoter, regulating the PIC formation, bears similarity to KMT2F’s binding to transcription initiation sites of RNA Pol II, we decided to explore this further. We hypothesized that the N terminal RRM (RNA recognition motif) of the KMT2F, which interacts with the WDR82, might be playing a similar role in the recruitment of the KMT2F to the rDNA locus through the WDR82 axis. To test this hypothesis, we cloned the fragment of the RRM spanning 74-172 amino acids in the SFB triple epitope vector, and ectopically expressed it in the cells. In our pull-down experiments with S-protein beads, the RRM showed association with the RNA Pol I, RRN3 and PAF-49 (Supplementary Figure S7C). As RRN3 is bound to active RNA Pol I only at the rDNA promoter, our results imply that the KMT2F may be recruited to the rDNA promoter, indirectly through WDR82.

### The catalytic activity of KMT2F is required for a stable PIC at the rDNA promoter

We have shown the detailed interaction between the KMT2F complex and various components of the RNA Pol I machinery in Figure 3. However, it remains unclear whether the direct physical interaction of KMT2F is responsible for RNA Pol I recruitment to rDNA or if this process depends on the catalytic activity of KMT2F. To investigate this, we conducted affinity pull down studies in cells ectopically expressing full length KMT2F or the catalytic dead mutant of KMT2F having a point mutation in the SET domain (KMT2F N1646A). Our S-protein pull-down assays revealed that KMT2F interacts with of RNA Pol I subunits as well as RRN3, and this interaction is unaffected with SET domain mutant KMT2F (Supplementary Figure S7D-E). Consistent with our results in S7C, these results show that the interaction of the RNA Pol I complex with the KMT2F is independent of its catalytic activity. As PIC formation is lost upon KMT2F knock down, we reasoned that maybe its catalytic activity is required to stabilize the PIC on the rDNA promoter. Indeed, when we performed ChIP of RNA Pol I and UBF, on these cells, in background of KMT2F shRNA, the catalytic dead mutant KMT2F could not sustain PIC formation, unlike the wild type KMT2F. These results indicated that H3K4me3 deposition by KMT2F is crucial for RNA Pol I PIC formation at rDNA promoter. Further, they also indicate that KMT2F recruits and stabilizes RNA Pol I PIC formation at rDNA promoter using its catalytic activity to promote transcription of 47S rRNA.

### KMT2A and KMT2F regulate rRNA transcription synergistically

Next, we asked, if KMT2F is able to promote PIC formation at rDNA promoter, what is the requirement of KMT2A in rRNA transcription? To answer this question, we overexpressed KMT2A alone, KMT2F alone or both KMT2Aand KMT2F. We observed that overexpression of KMT2A alone resulted in a modest 2-fold increase in rRNA transcripts, while overexpression of KMT2F alone showed 10-fold increase (Figure 6E, S7G). Surprisingly, co-expression of both proteins exhibited about 40-fold increase indicating that their activities are synergistic to rRNA transcription. Taken together, our results indicate that KMT2F modulates rRNA transcription via its KMT activity while KMT2A acts with KMT2F to augment the activity of RNA Pol I.

## Discussion

The H3K4me3 mark, deposited by the KMT2/COMPASS proteins, decorates the transcriptionally active promoters and is widely believed to be associated in activating RNA Pol II-mediated transcription (Lauberth et al., 2013). Here we show that the H3K4me3 mark deposited by KMT2F shapes the epigenetic landscape of the rDNA and promotes the PIC formation by RNA Pol I for active rRNA transcription. Our results highlights the similarities and differences in how KMT2 enzymes engages with the two RNA polymerases to promote transcription.

### KMT2 complexes interact with RNA Pol I

The three RNA polymerase I-III exist in homeostasis and if the functions of one are affected, they manifest in the transcriptional output of other two polymerases (Werner & Grohmann, 2011) (Vannini & Cramer, 2012) In order to elucidate a direct functional role of KMT2 complexes on the RNA Pol I machinery, and to rule out an indirect affect due to mis regulation of RNA Pol II by these complexes, we investigated the protein-protein interaction of COMPASS complex with RNA Pol I components. Our pull-down experiments demonstrated that WDR5 interacted with RNA Pol I subunits as well as with RRN3 and the SL1 complex, factors involved in transcription initiation, as well as with transcription elongation. In support of our findings, WDR5 was identified as a nucleolar protein in human proteome analysis and depletion of WDR5 has been reported to reduced rDNA transcription (Feng et al., 2010). Specifically, we show that WDR5 directly interacted with RPA34 and RPA49, peripheral subunits, specific to RNA Pol I. RPA34 and RPA49 form a functional heterodimer at RNA Pol I and have been implicated in RNA Pol I transcription initiation and elongation (Kuhn et al., 2007). Further, they exhibit functional similarity with the TFIIE and TFIIF modules, which regulate RNA Pol II initiation and elongation (Jeronimo & Robert, 2017)(Tsai et al., 2017)(Robinson et al., 2016). Consequently, it is plausible that KMTs may modulate RNA Pol I transcription through these interactions. Similarly, we explored the interaction of RNA Pol I specific subunits with KMT2A and KMT2F. Our endogenous immunoprecipitation experiments revealed that both the N- and C- subunits of KMT2A, as well as KMT2F, effectively precipitated the catalytic RNA Pol I transcriptional component RPA194, as well as UBF, and RRN3. We further mapped the RNA Pol I-interaction domain to RRM of KMT2F. Taken together, our results indicate that the regulation exerted by the KMT2 complexes involves physical proximity with the RNA Pol I complex.

### Unravelling the influence of KMT2 enzymes on epigenetic landscape of rDNA

The transcription process of rRNA genes necessitates a specialized RNA polymerase (RNA Pol I) and a distinct set of basal transcription factors (I. Grummt, 2003)(Moss et al., 2007). The basal transcription factors typically include UBF and the SL1 complex (consisting of TBP and TAFs), which play crucial roles in rDNA transcription. Our comprehensive mapping of RNA Pol I and UBF across the human rDNA locus was consistent with their specific binding patterns across the gene body and within the promoter region reported before. Minimal mapping of RNA Pol I was observed in the IGS region, consistent with previous studies (Herdman et al., 2017)(Mars et al., 2018)(Zentner et al., 2011). Earlier research has proposed that epigenetic mechanisms play a role in regulating the transcriptional activity of rRNA genes. Specifically, histone modifications emerge as pivotal elements in mediating the regulation of rDNA transcription (F. Yu et al., 2015). UBF exhibit a unique pattern of interaction with methylated H3K4, as its binding to rDNA locus is stimulated by H3K4me2 (Wu et al., 2017). Moreover, depletion of UBF results in decreased levels of H3K4me3 on the human rDNA promoter (Yu et al., 2015). These studied prompted the detailed global bioinformatic analysis of the distribution pattern of histone modifications on rDNA locus. The analysis revealed the distribution of six active histone modifications (H3K4me1, H3K4me2, H3K4me3, H3K9ac, H3K27ac and H3K36me3) and three repressive histone modifications (H3K9me1, H3K27me3 and H4K20me1) across the rDNA locus (Zentner et al., 2011). Given the pivotal role of KMT2 enzymes in regulating the distribution of H3K4 methylation marks across the genome, we were encouraged to unravel their direct functional significance on the regulation of rDNA locus. We choose KMT2A and KMT2F to conduct ChIP-seq analysis to delineate their distribution patterns on the rDNA locus. Our findings revealed the enrichment of both enzymes on the rDNA locus, with KMT2A exhibiting a preference for the intergenic region, while KMT2F demonstrated a nearly uniform distribution pattern in the RNA Pol I transcribed region. When we performed ChIP-seq analysis for H3K4me2 and H3K4me3, it corelated more closely with KMT2F (Figure S2). To delve deeper into the mechanistic details of how these two proteins regulate the rDNA locus, we tried to generate CRISPR-mediated knockouts of KMT2F. However, as CRISPR-mediated knock outs cells of KMT2F were not viable, we were limited to use shRNA-mediated knockdown of KMT2F. Our observations revealed active regulation of active signatures (H3K4me2 and H3K4me3) following the loss of this protein. KMT2F depletion exhibited decrease in H3K4me2, H3K4me3, H3K9ac marks and corresponding increase in H3K9me3 mark on promoter and transcribed unit. Therefore, our results reveal that KMT2F-mediated H3K4me3 deposition not only acts directly by regulating RNA Pol I PIC formation but also indirectly, by shaping the epigenetic landscape of the rDNA.

### Drawing comparison between RNA Pol I or RNA Pol II transcription processes and role of KMT2F therein

Although the eukaryotic RNA polymerases differ from each other by performing a vast array of different functions, but they share structural similarities, in terms of their catalytic core and in the functional steps of the basic transcription process. For instance, despite having 14 subunits (RNA Pol I) and 12 subunits (RNA Pol II), both RNA polymerase still share 7 common subunits. Further, functional similarities, like those exemplified by the heterodimeric complex of RNA Pol I (RPA49-RPA34) functionally resembling the general transcription factor TFIIE-TFIIF modules of RNA Polymerase II, are observed. Similarly, the transcription factor TFIIB plays an analogous role as played by RNA Pol I associated factor RRN3 in transcription initiation (Kuhn et.al; 2007). There are examples of factors which resemble both structurally and functionally in the other processes of the transcription like in the elongation, regulation etc (Zhang et al., 2008). We observe many similarities in how KMT2F engages with RNA Pol I comparable to its engagement with RNA Pol II. Although the KMT2F lacks the direct DNA binding domains, but it interacts with WDR82 or CFP1, which targets it to the chromatin through the RNA Pol II or through the CXXC of CFP1. Looking at the broad similarities in the two processes, and our results showing that KMT2F interacts with RNA Pol I components through its RRM domain, we hypothesize a parallel recruitment mechanism for KMT2F to the rDNA locus.

KMT2F plays a critical role in the global deposition of H3K4me3 marks, which have been widely observed to support transcriptional activation of RNA Pol II regulated genes. H3K4me3 contributes to the formation of the RNA Pol II PIC at active promoters. The TAF3 subunit of TFIID interacts with H3K4me3 to facilitate the recruitment of the TFIID complex, consisting of the TATA-binding protein (TBP) and nearly 14 TBP associated factors (TAFs) (Lauberth et al., 2013). Additionally, H3K4me3 mark can recruit chromatin remodelers, which maintain an open chromatin state at the gene promoters (H. Wang & Helin, 2024). However, recent reports suggest that H3K4me3 is crucial for RNA Polymerase II pause release (Hu et al., 2023)(H. Wang et al., 2023). Although there is growing evidence supporting the role of H3K4me3 in regulating transcription by RNA Pol II, its role in modulating rDNA transcription by RNA Pol I remains surprisingly unexplored. Our findings indicate that H3K4me3, deposited by KMT2F, is crucial for pre-initiation complex formation on rDNA promoters.

Interestingly, KMT2F physically interacts with RNA Pol I but needs its KMT activity to stabilize the RNA Pol I PIC on the promoter. In contrast to the similarities outlined above between the two polymerase processes, these findings present a divergence; as depletion of KMT2F, or mutations in its SET domain, do not affect RNA Pol II recruitment to the chromatin. However, our results reveal that loss of KMT2F catalytic activity disrupts RNA Pol I recruitment to rDNA. Many factors may contribute to these differences. For instance, recycling and transcription re-initiation by RNA Pol I, dependence of PIC factors like UBF on H3K4 methylation, organization and accessibility of rDNA chromatin, may individually or together necessitate the presence of H3K4me3 on rDNA promoter for PIC formation. This suggests a possible co-dependence of RNA Pol I on KMT2F for proper recruitment at the rDNA, highlighting distinct roles that KMTs play in different transcriptional contexts.

### Impact of KMT2F-mediated rDNA transcription on disease biology

Heterozygous loss-of-function variations in KMT2F have been associated with various pathologies, including childhood speech ataxia (Eising et al., 2019), early-onset epilepsy (X. Yu et al., 2019) and severe developmental disorders (Singh et al., 2016). However, the most notable association of KMT2F lies in the development of schizophrenia. Patients diagnosed with schizophrenia have shown increased transcriptional activity of rDNA in both brain tissue and lymphocytes. This heightened activity may arise from elevated rDNA content in the genomes of individuals with schizophrenia compared to healthy controls (Krzyzanowska et al., 2015)(Veiko et al., 2003). This proposed interconnection among schizophrenia, KMT2F, and rDNA transcription underscores the importance of further investigating the role of KMT2F in rDNA transcription. Taken together these studies indicate the necessity for deeper investigation into how KMT2F may deregulate the transcription of the rDNA locus in disease condition, while our study firmly establishes its regulatory role in RNA Pol I transcription.

## Acknowledgements

We thank Ingrid Grummit for providing the RPA43 and RPA49 constructs. We are thankful Z.U. Zargar for assistance with initial ChIP experiments, K. Malik for assistance with siRNA blots and Ajay Mahato, Asmita Gupta and Dharitri Saikia for their contributions to ChIP-seq analysis.

## Author Contributions

K.A.L. conducted all experiments and ChIP-seq analysis, except ones noted below. A.K. conducted experiments presented in Fig. 1A, 3B, 3D, 4A, 4D-G, S1, S4B-F, S5. G.R. performed experiments presented in Fig. 3C, 3E and Fig. S4A. S.T., A.K., G.R., and K.A.L. designed the experiments. S.T. and K.A.L. wrote the manuscript.

## Material and Methods

### Cloning and site-directed mutagenesis

The cDNA sequences of RPA12, RPA34, RRN3, TAF63, and TAF48 were PCR amplified from the cDNA synthesized from total RNA. Subsequently, the amplified cDNA was cloned into the TOPO vector using the TOPO TA Cloning Kit (Invitrogen), and then subcloned into pGEX4T1 to generate GST-tagged constructs. GST RPA43 was generated by cloning the PCR product amplified from TOPO RPA43 (a kind gift from Ingrid Grummt) into pGEX4T1. GST-KMT2A-SET (GST-MLL-D3, Ali et al., 2017) and GST-RPA49 (a kind gift from Ingrid Grummt, Yuan et al.,2002) has been described before. RRM was PCR amplified from PCDNA-SFB-KMT2F vector and then sub cloned in the Xho1 site of the PCDNA-SFB vector.

### Cell culture, stable cell line generation and transfections

HEK-293 (human embryonic kidney), HeLa (Cervical Cancer cell line), MCF-7 (Breast Cancer cell line), U-2OS (human osteosarcoma), and IMR-90 tert (human lung fibroblast) cells were cultured in DMEM (Gibco), supplemented with 10% (v/v) fetal bovine serum (FBS), 1% (v/v) GlutaMAX, and 100 U/ml penicillin-streptomycin. Authentication of cell lines was conducted via STR profiling (Life Technologies). The cells were maintained at 37°C in a humidified atmosphere with 5% CO2. Generation of full-length and mutant KMT2A and KMT2F cell lines have been described previously (Ali et al., 2014; Malik et al., 2023). Similar methods were used in the generation of KMT2F full length and mutant cell lines in the HEK-293 cells (Malik et al., 2023). To deplete the endogenous proteins, RNA interference (RNAi) was 24 employed targeting KMT2A, KMT2B, KMT2C, KMT2F, and WDR5 genes, as reported previously (Ali et al., 2014)(Chinchole et al., 2022; Malik et al., 2023). Briefly, siRNA duplexes were designed and transfected into the cells using Oligofectamine (Thermo Fisher Scientific) as the transfecting agent. A siRNA sequence targeting the firefly luciferase gene was used as a control. KMT2F shRNAs were transiently transfected into HEK-293 cells using PEI, as described previously (Malik et al., 2023). Cells underwent two rounds of transfections at a time interval of 24 hours and were collected after 72 hours.

### Protein expression and purification

All GST-tagged proteins were expressed using either BL 21 (DE3) or Rosetta Gami DE3 *Escherichia coli* strain (Novagen). Protein expression was induced by adding 0.1 mM Isopropyl β-D-1-thiogalactopyranoside (IPTG) for 6 hours at 18°C when the culture reached 0.6 OD. The cells were then pelleted and lysed in lysis buffer (50 mM Tris pH 7.5, 150 mM NaCl, 0.5% NP-40, and 1 mM PMSF), followed by incubation with glutathione agarose beads (Sigma) at 4°C for 2-3 hours. Subsequently, the beads underwent washing with ice-cold lysis buffer 3–4 times. Protein concentration was determined via SDS-PAGE followed by CBB staining.

### Co-immunoprecipitation, GST Affinity pull-downs and immunoblot analysis

Mammalian cell lysates were prepared by either performing whole cell lysis using NETN (100mM NaCl, 20 mM Tris-HCl pH 8, 0.5 mM EDTA, and 0.5% Nonidet P-40) or specifically conducting nuclear lysis (Dignam, Lebovitz, and Roeder 1983). Cell/nuclear lysates were treated with a cocktail of protease inhibitors (Sigma) and was used for immunoprecipitation/pull-down experiments. The pre-cleared lysate was incubated with 1microg of IgG, KMT2A or KMT2F antibodies respectively or with S protein beads for SFB tagged proteins at 40C O/N. Following incubation with antibody, the cell lysate for endogenous pulldowns was incubated 25 with 20ml of 50% slurry of Protein A beads for 1 hour at 40C. The pelleted Protein A/ S protein beads were washed 3 times with IP wash buffer (15mM TrisCl pH 7.5, 100mM KCL, 0.02%NP-40) and boiled in Laemmli buffer. The samples were run on a SDS-PAGE followed by immunoblotting. For pull-down experiments involving GST-tagged proteins, an equal amount of purified bead-bound proteins were incubated with HeLa or HeLa spinner whole-cell or nuclear extracts for 2-3 hours, followed by washing with IP wash buffer. The boiled protein extracts were separated by SDS-PAGE and transferred to either nitrocellulose (Amersham, 10600003) or PVDF (Amersham, 10600023) membranes. Endogenous KMT2A-C subunit (A300-374A, Bethyl Labs), KMT2A-N subunit (A300-086A, Bethyl Labs), and KMT2F (A300-289A, Bethyl Labs) were pulled down and immunoblotted with RPA194 (SC-48385, Santa Cruz), UBF (SC-13125, Santa Cruz), RRN3 (Ab-112052, Abcam), and KMT2A or KMT2F antibodies mentioned above. Similarly, SFB pull downs were probed with RPA-49 (HPA022-527, Sigma), RPA194 (SC-48385, Santa Cruz), RRN3 (Ab-112052, Abcam) or with FLAG (F7425, Sigma). Blots were scanned using an Odyssey infrared imager (LI-COR) or ImageQuant LAS 500.

### *in-vitro* interaction studies

Thrombin enzyme (Sigma) was utilized to cleave the GST tag in purified GST-WDR5 at 22°C for 18 hours. The intact WDR5 protein post-cleavage was retrieved and employed for *in-vitro* interaction studies. Subsequently, bacterially purified GST tag proteins were subjected to incubation with purified WDR5 (500 ng each) in interaction buffer (IP dilution buffer: 50 mM TRIS pH 7.4, 150 mM NaCl, 0.1 mM EDTA, 0.1% NP40, 100 μg/ml BSA also containing 200 mM KCl) for a duration of 12 hours. Beads were collected by centrifugation, followed by thorough washing with IP wash buffer. Finally, Western blotting was employed to identify bound protein-complexes.

### Immunofluorescence imaging

Cells (U-2OS, MCF-7, HeLa, and IMR90-tert) were cultured on poly lysine-coated glass cover slips. Different fixation protocols were employed to detect the localization of endogenous KMT2A and KMT2F proteins. For KMT2A, cells were fixed with pre-chilled acetone for 4 minutes at - 20°C, while KMT2F fixation was achieved with pre-chilled methanol and acetic acid (1:100). Subsequently, fixed cells were blocked with one percent BSA for 30 minutes at room temperature. The cells were then incubated overnight at 4°C with antibodies against KMT2A-C (1:100) (A300-374A, Bethyl Labs), KMT2F (1:100) (A300-289A, Bethyl Labs), and B23/nucleophosmin (1:150) (B0556, Sigma), used as a marker for the nucleolus. Following primary antibody incubation, the cells were washed and incubated with Alexa 488 (1:1000, Thermo Fisher Scientific A11034), Alexa 633 (1:500, Thermo Fisher Scientific A21050), and/or Alexa 594 (1:500, Thermo Fisher Scientific A11030) conjugated anti-rabbit or anti-mouse secondary antibodies at room temperature for 1 hour. The samples were mounted in VECTASHIELD Mounting Medium (Vector Laboratories-H1200) with 4-6-diamidino-2-phenylindole (DAPI) to stain the DNA. Images were acquired using a ZEISS LSM 900 inverted confocal microscope with a 63×/1.4 oil immersion lens.

### RNA isolation and qRT-PCR

Total RNA isolation and cDNA preparation was done as described (Malik et al., 2023). Before cDNA synthesis, the RNA was treated with TURBO DNase (Thermo Fisher Scientific) for 30 minutes at 37°C to eliminate genomic DNA contamination. To ensure RNA purity, a no-enzyme RNA amplification step was conducted prior to cDNA synthesis. After confirming the absence of DNA contamination, cDNA synthesis was carried out using the SuperScript III Reverse Transcriptase kit (Thermo Fisher Scientific) following the manufacturer’s instructions. RT-qPCR analysis was conducted using either the 7500 Real-Time PCR (Applied Biosystems), QuantStudio 5 Real-Time PCR System (Applied Biosystems), or Bio-Rad (CFX-maestro) platforms, employing the DyNAmo ColorFlash SYBR Green qPCR kit (Thermo Fisher Scientific). Transcript levels were quantified using the 2-ΔΔCt method (Livak and Schmittgen 2001). Primer sequences are provided in Supplementary Table S1.

### [32P] metabolic labeling and nascent ribosomal RNA quantification

Cells were incubated in phosphate free media containing 10% FBS for 2 h followed by incubation with DMEM containing 0.1 mCi/ ml of [32P] orthophosphate for 1 h. Labelled RNA was extracted by TRIzol (Ambion) reagent. RNA was quantified with NanoDrop, divided into two portions for each sample from different treatments and one portion was loaded on 1.2% agarose gel and ran at 60 Volt/centimeter for 5 h. Gel was further dried at 8^0^C for 1h. RNA was visualized by autoradiography (Typhoon FLA-9500, GE) and densitometry quantification was done by Phosphorimaging. 45S ribosomal RNA levels were quantified between the control and treatment by normalizing 45S RNA levels with 28S ribosomal RNA levels. To visualize 28S rRNA, second portion of the sample was run on denaturing agarose gel containing nine percent formaldhyde and visualized after staining with ethidium bromide.

### Chromatin immunoprecipitation

Chromatin immunoprecipitation (ChIP) experiments were performed as previously described by (Malik et al. 2023) (Zargar, Kimidi, and Tyagi 2018). using the following antibodies: KMT2A-C (A300-374A, Bethyl Labs or in-house as described in (Chinchole, Lone, and Tyagi 2022) KMT2F (A300-289A, Bethyl Labs), H3K4me2 (ab32356, Abcam); H3K4me3 (ab8580, Abcam or 07-473 Millipore); H3 (ab1791, Abcam); H3K9me3 (ab8898, Abcam); H3K9Ac (ab4441, Abcam); RPA194 (SC-48385, Santa cruz); UBF (SC-13125, Santa cruz); RRN3 (Ab-112052, Abcam); and TAF1C (A303-698A, Bethyl labs);. The relative occupancy or percent input of the immunoprecipitated protein at ribosomal DNA locus was estimated by RT-qPCR as follows: 100 × 2(Ct Input–Ct IP), where Ct Input and Ct IP are mean threshold cycles of RT-qPCR on DNA samples from input and specific immunoprecipitations, respectively. To measure fold over control, fold change over the ChIP values obtained in the control cells was used. Primer sequences are provided in Supplementary Table S1.

### ChIP seq and analysis

The ChIP DNA (10 ng) of KMT2A and KMT2F from HEK-293 cells was utilized to construct ChIP-sequencing libraries employing the NEB Next Ultra II DNA Library preparation kit. Subsequently, paired-end sequencing (2 × 150 bp) was carried out on the Illumina Nextseq 2000 platform, utilizing the sequencing services provided by the CDFD sequencing facility (National Genomics Core (NGC), CDFD). The ChIP-seq datasets of KMT2A and KMT2F, as well as previously published ChIP-seq datasets of H3K4me2 (GSM5954234), H3K4me3 (GSM5954235), Input for H3K4me2 and H3K4me3 (GSM5954242) (Liu et al. 2022) and UBF (SRX035783), RNA Pol I (SRX035784) Input for UBF and Pol I (SRX035785) (Zentner et al. 2011) for WDR82 (GSE186758) (Park et al., 2022) and for WDR5 (GSE60897) (Thomas et al., 2015) were analyzed to assess their binding on human rDNA, using a pipeline previously described for rDNA (George, Pimkin, and Paralkar 2023). Briefly, the quality of raw sequence data from experimental (IP) and input DNA samples was assessed using FastQC version 0.11.4 (https://www.bioinformatics.babraham.ac.uk/projects/fastqc/). Adapter removal from raw sequencing read pairs was performed using TrimGalore (v0.6.7) (https://github.com/FelixKrueger/TrimGalore) or Trimmomatic. The trimmed sequencing reads were then mapped to the customized human reference genome (hg38-rDNA) using indexed Bowtie2 (version 2.3.2). SAM files obtained after alignment were converted to BAM format using Samtools (version 1.9). Sorting and indexing of BAM files was done using Samtools (version 1.9). Visualization tracks were created using DeepTools (BamCoverage). Subsequently, BigWig (BW) files generated for input and experimental data (IP) post-BamCoverage were compared using BW Compare(https://deeptools.readthedocs.io/en/develop/content/tools/bigwigCompare.html). The individual BW files for each dataset, generated after comparison (displayed in log2 scale), were subsequently visualized using IGV (Integrated Genomic Viewer).

## References

Ali, A., Veeranki, S. N., & Tyagi, S. (2014). A SET-domain-independent role of WRAD complex in cell-cycle regulatory function of mixed lineage leukemia. Nucleic Acids Research, 42(12), 7611–7624. 10.1093/nar/gku458

Bochyńska, A., Lüscher-Firzlaff, J., & Lüscher, B. (2018). Modes of interaction of KMT2 histone H3 lysine 4 methyltransferase/COMPASS complexes with chromatin. In Cells (Vol. 7, Issue 3). MDPI. 10.3390/cells7030017

Brown, David A et al. 2017. “The SET1 Complex Selects Actively Transcribed Target Genes via Multivalent Interaction with CpG Island Chromatin.” Cell reports 20(10): 2313–27. http://www.ncbi.nlm.nih.gov/pubmed/28877467.

Caslini, C., Alarcòn, A. S., Hess, J. L., Tanaka, R., Murti, K. G., & Biondi, A. (2000). The amino terminus targets the mixed lineage leukemia (MLL) protein to the nucleolus, nuclear matrix and mitotic chromosomal scaffolds. Leukemia, 14(11), 1898–1908. 10.1038/sj.leu.2401933

Chinchole, A., Lone, K. A., & Tyagi, S. (2022). MLL regulates the actin cytoskeleton and cell migration by stabilising Rho GTPases via the expression of RhoGDI1. Journal of Cell Science, 135(20). 10.1242/JCS.260042/VIDEO-3

Comai, L., Zomerdijk, J. C. B. M., Beckmann, H., Zhou, S., Admon, A., & Tjian, R. (1994). Reconstitution of Transcription Factor SL1: Exclusive Binding of TBP by SL1 or TFIID Subunits. Science, 266(5193), 1966–1972. 10.1126/SCIENCE.7801123

Denissov, S., Hofemeister, H., Marks, H., Kranz, A., Ciotta, G., Singh, S., Anastassiadis, K., Stunnenberg, H. G., & Stewart, A. F. (2014). Mll2 is required for H3K4 trimethylation on bivalent promoters in embryonic stem cells, whereas Mll1 is redundant. Development (Cambridge), 141(3), 526–537. 10.1242/dev.102681

Dignam, J. D., R. M. Lebovitz, and R. G. Roeder. 1983. “Accurate Transcription Initiation by RNA Polymerase II in a Soluble Extract from Isolated Mammalian Nuclei.” Nucleic Acids Research 11(5): 1475–89. https://academic.oup.com/nar/article-lookup/doi/10.1093/nar/11.5.1475.

Dixon, J., Trainor, P., & Dixon, M. J. (2007). Treacher Collins syndrome. Orthodontics and Craniofacial Research, 10(2), 88–95. 10.1111/j.1601-6343.2007.00388.x

Eberhard, D., Tora, L., Egly, J. marc, & Grummt, I. (1993). A TBP-containing multiprotein complex (TIF-IB) mediates transcription specificity of murine RNA polymerase I. Nucleic Acids Research, 21(18), 4180. 10.1093/NAR/21.18.4180

Eising, E., Carrion-Castillo, A., Vino, A., Strand, E. A., Jakielski, K. J., Scerri, T. S., Hildebrand, M. S., Webster, R., Ma, A., Mazoyer, B., Francks, C., Bahlo, M., Scheffer, I. E., Morgan, A. T., Shriberg, L. D., & Fisher, S. E. (2019). A set of regulatory genes co-expressed in embryonic human brain is implicated in disrupted speech development. Molecular Psychiatry, 24(7), 1065–1078. 10.1038/s41380-018-0020-x

Ernst, P., & Vakoc, C. R. (2012). WRAD: Enabler of the SET1-family of H3K4 methyltransferases. Briefings in Functional Genomics, 11(3), 217–226. 10.1093/bfgp/els017

Feng, W., Yonezawa, M., Ye, J., Jenuwein, T., & Grummt, I. (2010). PHF8 activates transcription of rRNA genes through H3K4me3 binding and H3K9me1/2 demethylation. Nature Structural and Molecular Biology, 17(4), 445–450. 10.1038/nsmb.1778

Franks, Tobias M et al. 2017. “Nup98 Recruits the Wdr82-Set1A/COMPASS Complex to Promoters to Regulate H3K4 Trimethylation in Hematopoietic Progenitor Cells.” Genes & development 31(22): 2222–34. http://www.ncbi.nlm.nih.gov/pubmed/29269482.

Friedrich, J. K., Panov, K. I., Cabart, P., Russell, J., & Zomerdijk, J. C. B. M. (2005). TBP-TAF complex SL1 directs RNA polymerase I pre-initiation complex formation and stabilizes upstream binding factor at the rDNA promoter. Journal of Biological Chemistry, 280(33), 29551–29558. 10.1074/jbc.M501595200

Ganapathi, K. A., Austin, K. M., Lee, C. S., Dias, A., Malsch, M. M., Reed, R., & Shimamura, A. (2007). The human Shwachman-Diamond syndrome protein, SBDS, associates with ribosomal RNA. Blood, 110(5), 1458–1465. 10.1182/blood-2007-02-075184

George, S. S., Pimkin, M., & Paralkar, V. R. (2023). Construction and validation of customized genomes for human and mouse ribosomal DNA mapping. Journal of Biological Chemistry, 299(6). 10.1016/j.jbc.2023.104766

Goodfellow, S. J., & Zomerdijk, J. C. B. M. (2013). Basic mechanisms in RNA polymerase I transcription of the ribosomal RNA genes. Sub-Cellular Biochemistry, 61, 211–236. 10.1007/978-94-007-4525-4_10

Gorski, J. J., Pathak, S., Panov, K., Kasciukovic, T., Panova, T., Russell, J., & Zomerdijk, J. C. B. M. (2007). A novel TBP-associated factor of SL1 functions in RNA polymerase I transcription. EMBO Journal, 26(6), 1560–1568. 10.1038/sj.emboj.7601601

Grummt, I. (1982). Nucleotide sequence requirements for specific initiation of transcription by RNA polymerase I (rDNA promoter/cell-free transcription/deletion mutants/upstream sequences). In Proc. NatL Acad. Sci. USA (Vol. 79).

Grummt, I. (2003). Life on a planet of its own: regulation of RNA polymerase I transcription in the nucleolus. Genes & Development, 17(14), 1691–1702. 10.1101/GAD.1098503R

Grummt, L., Rosenbauer, H., Niedermeyer, L., Maier, U., & Dhrlein, A. (1986). A Repeated 18 bp Sequence Motif in the Mouse rDNA Spacer Mediates Binding of a Nuclear Factor and Transcription Termination. In Cell (Vol. 45).

Hannan, K. M., Sanij, E., Rothblum, L. I., Hannan, R. D., & Pearson, R. B. (2013). Dysregulation of RNA polymerase I transcription during disease. In Biochimica et Biophysica Acta - Gene Regulatory Mechanisms (Vol. 1829, Issues 3–4, pp. 342–360). 10.1016/j.bbagrm.2012.10.014

Heiss, N S et al. 1998. “X-Linked Dyskeratosis Congenita Is Caused by Mutations in a Highly Conserved Gene with Putative Nucleolar Functions.” Nature genetics 19(1): 32–38. http://www.ncbi.nlm.nih.gov/pubmed/9590285.

Herdman, C., Mars, J. C., Stefanovsky, V. Y., Tremblay, M. G., Sabourin-Felix, M., Lindsay, H., Robinson, M. D., & Moss, T. (2017). A unique enhancer boundary complex on the mouse ribosomal RNA genes persists after loss of Rrn3 or UBF and the inactivation of RNA polymerase I transcription. PLoS Genetics, 13(7). 10.1371/journal.pgen.1006899

Hu, S., Song, A., Peng, L., Tang, N., Qiao, Z., Wang, Z., Lan, F., & Chen, F. X. (2023). H3K4me2/3 modulate the stability of RNA polymerase II pausing. In Cell Research (Vol. 33, Issue 5, pp. 403–406). Springer Nature. 10.1038/s41422-023-00794-3

Jeronimo, C., & Robert, F. (2017). The Mediator Complex: At the Nexus of RNA Polymerase II Transcription. In Trends in Cell Biology (Vol. 27, Issue 10, pp. 765–783). Elsevier Ltd. 10.1016/j.tcb.2017.07.001

Karole, A. M., Chodisetty, S., Ali, A., Kumari, N., & Tyagi, S. (2018). Novel sub-cellular localizations and intra-molecular interactions may define new functions of Mixed Lineage Leukemia protein. Cell Cycle, 17(24), 2684–2696. 10.1080/15384101.2018.1553338

Krzyzanowska, M., Steiner, J., Brisch, R., Mawrin, C., Busse, S., Braun, K., Jankowski, Z., Bernstein, H. G., Bogerts, B., & Gos, T. (2015). Ribosomal DNA transcription in dorsal raphe nucleus neurons is increased in residual schizophrenia compared to depressed patients with affective disorders. Psychiatry Research, 230(2), 233–241. 10.1016/j.psychres.2015.08.045

Kuhn, A., & Grummt, I. (1987). A novel promoter in the mouse rDNA spacer is active in vivo and in vitro. In The EMBO Journal (Vol. 6, Issue 11).

Kuhn, C. D., Geiger, S. R., Baumli, S., Gartmann, M., Gerber, J., Jennebach, S., Mielke, T., Tschochner, H., Beckmann, R., & Cramer, P. (2007). Functional Architecture of RNA Polymerase I. Cell, 131(7), 1260– 1272. 10.1016/j.cell.2007.10.051

Labhart, P, and R H Reeder. 1986. “Characterization of Three Sites of RNA 3’ End Formation in the Xenopus Ribosomal Gene Spacer.” Cell 45(3): 431–43. http://www.ncbi.nlm.nih.gov/pubmed/3453104.

Lauberth, S. M., Nakayama, T., Wu, X., Ferris, A. L., Tang, Z., Hughes, S. H., & Roeder, R. G. (2013). H3K4me3 interactions with TAF3 regulate preinitiation complex assembly and selective gene activation. Cell, 152(5), 1021–1036. 10.1016/j.cell.2013.01.052

Learned, R. M., Cordes, S., & Tjian, R. (1985). Purification and Characterization of a Transcription Factor That Confers Promoter Specificity to Human RNA Polymerase I. In MOLECULAR AND CELLULAR BIOLOGY (Vol. 5, Issue 6).

Lee, J.-H., & Skalnik, D. G. (2008). Wdr82 Is a C-Terminal Domain-Binding Protein That Recruits the Setd1A Histone H3-Lys4 Methyltransferase Complex to Transcription Start Sites of Transcribed Human Genes. Molecular and Cellular Biology, 28(2), 609–618. 10.1128/mcb.01356-07

Li, Y., Zhao, L., Zhang, Y., Wu, P., Xu, Y., Mencius, J., Zheng, Y., Wang, X., Xu, W., Huang, N., Ye, X., Lei, M., Shi, P., Tian, C., Peng, C., Li, G., Liu, Z., Quan, S., & Chen, Y. (2022). Structural basis for product specificities of MLL family methyltransferases. Molecular Cell, 82(20), 3810–3825.e8. 10.1016/j.molcel.2022.08.022

Liu, Hongsen et al. 2022. “The Non-Specific Lethal (NSL) Histone Acetyltransferase Complex Transcriptionally Regulates Yin Yang 1-Mediated Cell Proliferation in Human Cells.” International Journal of Molecular Sciences 23(7): 3801. https://www.mdpi.com/1422-0067/23/7/3801.

Liu, X., Wang, C., Liu, W., Li, J., Li, C., Kou, X., Chen, J., Zhao, Y., Gao, H., Wang, H., Zhang, Y., Gao, Y., & Gao, S. (2016). Distinct features of H3K4me3 and H3K27me3 chromatin domains in pre-implantation embryos. Nature, 537(7621), 558–562. 10.1038/nature19362

Livak, K J, and T D Schmittgen. 2001. “Analysis of Relative Gene Expression Data Using Real-Time Quantitative PCR and the 2(-Delta Delta C(T)) Method.” Methods (San Diego, Calif.) 25(4): 402–8. http://www.ncbi.nlm.nih.gov/pubmed/11846609

Malik, K. K., Sridhara, S. C., Lone, K. A., Katariya, P. D., Pulimamidi, D., & Tyagi, S. (2023). MLL methyltransferases regulate H3K4 methylation to ensure CENP-A assembly at human centromeres. PLOS Biology, 21(6), e3002161. 10.1371/journal.pbio.3002161

Mars, J. C., Sabourin-Felix, M., Tremblay, M. G., & Moss, T. (2018). A deconvolution protocol for ChIP-seq reveals analogous enhancer structures on the mouse and human ribosomal RNA genes. G3: Genes, Genomes, Genetics, 8(1), 303–314. 10.1534/g3.117.300225

McStay, B., & Grummt, I. (2008). The epigenetics of rRNA genes: From molecular to chromosome biology. In Annual Review of Cell and Developmental Biology (Vol. 24, pp. 131–157). 10.1146/annurev.cellbio.24.110707.175259

Moss, T., Langlois, F., Gagnon-Kugler, T., & Stefanovsky, V. (2007). A housekeeper with power of attorney: The rRNA genes in ribosome biogenesis. In Cellular and Molecular Life Sciences (Vol. 64, Issue 1, pp. 29–49). 10.1007/s00018-006-6278-1

Narla, A., & Ebert, B. L. (2010). Ribosomopathies: Human disorders of ribosome dysfunction. In Blood (Vol. 115, Issue 16, pp. 3196–3205). American Society of Hematology. 10.1182/blood-2009-10-178129

Nguyen, L. X. T., & Mitchell, B. S. (2013). Akt activation enhances ribosomal RNA synthesis through casein kinase II and TIF-IA. Proceedings of the National Academy of Sciences of the United States of America, 110(51), 20681–20686. 10.1073/pnas.1313097110

Panov, K. I., Friedrich, J. K., Russell, J., & Zomerdijk, J. C. B. M. (2006). UBF activates RNA polymerase I transcription by stimulating promoter escape. EMBO Journal, 25(14), 3310–3322. 10.1038/sj.emboj.7601221

Park, K., Zhong, J., Jang, J. S., Kim, J., Kim, H. J., Lee, J. H., & Kim, J. (2022). ZWC complex-mediated SPT5 phosphorylation suppresses divergent antisense RNA transcription at active gene promoters. Nucleic Acids Research, 50(7), 3835–3851. 10.1093/nar/gkac193

Robinson, P. J., Trnka, M. J., Bushnell, D. A., Davis, R. E., Mattei, P. J., Burlingame, A. L., & Kornberg, R. D. (2016). Structure of a Complete Mediator-RNA Polymerase II Pre-Initiation Complex. Cell, 166(6), 1411–1422.e16. 10.1016/j.cell.2016.08.050

Sankar, A., Lerdrup, M., Manaf, A., Johansen, J. V., Gonzalez, J. M., Borup, R., Blanshard, R., Klungland, A., Hansen, K., Andersen, C. Y., Dahl, J. A., Helin, K., & Hoffmann, E. R. (2020). KDM4A regulates the maternal-to-zygotic transition by protecting broad H3K4me3 domains from H3K9me3 invasion in oocytes. Nature Cell Biology, 22(4), 380–388. 10.1038/s41556-020-0494-z

Sharifi, S., & Bierhoff, H. (2018). Annual Review of Biochemistry Regulation of RNA Polymerase I Transcription in Development, Disease, and Aging. 10.1146/annurev-biochem

Shilatifard, A. (2012). The COMPASS family of histone H3K4 methylases: Mechanisms of regulation in development and disease pathogenesis. Annual Review of Biochemistry, 81, 65–95. 10.1146/annurev-biochem-051710-134100

Singh, T., Kurki, M. I., Curtis, D., Purcell, S. M., Crooks, L., McRae, J., Suvisaari, J., Chheda, H., Blackwood, D., Breen, G., Pietilinen, O., Gerety, S. S., Ayub, M., Blyth, M., Cole, T., Collier, D., Coomber, E. L., Craddock, N., Daly, M. J., … Barrett, J. C. (2016). Rare loss-of-function variants in SETD1A are associated with schizophrenia and developmental disorders. Nature Neuroscience, 19(4), 571–577. 10.1038/nn.4267

Sollner-Webb, B., Anne, J., Wilkinson, K., Roan, J., & Reedert, R. H. (1983). Nested Control Regions Promote Xenopus Ribosomal RNA Synthesis by RNA Polymerase I. In Cell (Vol. 35).

Stefanovsky, V., Langlois, F., Gagnon-Kugler, T., Rothblum, L. I., & Moss, T. (2006). Growth factor signaling regulates elongation of RNA polymerase I transcription in mammals via UBF phosphorylation and r-chromatin remodeling. Molecular Cell, 21(5), 629–639. 10.1016/j.molcel.2006.01.023

Stults, D. M., Killen, M. W., Pierce, H. H., & Pierce, A. J. (2008). Genomic architecture and inheritance of human ribosomal RNA gene clusters. Genome Research, 18(1), 13–18. 10.1101/gr.6858507

Sugeedha, J., Gautam, J., & Tyagi, S. (2021). SET1/MLL family of proteins: functions beyond histone methylation. In Epigenetics (Vol. 16, Issue 5, pp. 469–487). Bellwether Publishing, Ltd. 10.1080/15592294.2020.1809873

Thomas, L. R., Wang, Q., Grieb, B. C., Phan, J., Foshage, A. M., Sun, Q., Olejniczak, E. T., Clark, T., Dey, S., Lorey, S., Alicie, B., Howard, G. C., Cawthon, B., Ess, K. C., Eischen, C. M., Zhao, Z., Fesik, S. W., & Tansey, W. P. (2015). Interaction with WDR5 promotes target gene recognition and tumorigenesis by MYC. Molecular Cell, 58(3), 440–452. 10.1016/j.molcel.2015.02.028

Tsai, K. L., Yu, X., Gopalan, S., Chao, T. C., Zhang, Y., Florens, L., Washburn, M. P., Murakami, K., Conaway, R. C., Conaway, J. W., & Asturias, F. J. (2017). Mediator structure and rearrangements required for holoenzyme formation. Nature, 544(7649), 196–201. 10.1038/nature21393

Tyagi, S., Chabes, A. L., Wysocka, J., & Herr, W. (2007). E2F Activation of S Phase Promoters via Association with HCF-1 and the MLL Family of Histone H3K4 Methyltransferases. Molecular Cell, 27(1), 107–119. 10.1016/j.molcel.2007.05.030

Vannini, A., & Cramer, P. (2012). Conservation between the RNA Polymerase I, II, and III Transcription Initiation Machineries. In Molecular Cell (Vol. 45, Issue 4, pp. 439–446). 10.1016/j.molcel.2012.01.023

Veiko, N. N., Egolina, N. A., Radzivil, G. G., Nurbaev, S. D., Kosyakova, N. V., Shubaeva, N. O., & Lyapunova, N. A. (2003). Quantitation of Repetitive Sequences in Human Genomic DNA and Detection of an Elevated Ribosomal Repeat Copy Number in Schizophrenia: The Results of Molecular and Cytogenetic Analyses. Molecular Biology, 37(3), 349–357. 10.1023/A:1024274924381/METRICS

Viktorovskaya, O. V., & Schneider, D. A. (2015). Functional divergence of eukaryotic RNA polymerases: Unique properties of RNA polymerase I suit its cellular role. In Gene (Vol. 556, Issue 1, pp. 19–26). Elsevier B.V. 10.1016/j.gene.2014.10.035

Wang, H., Fan, Z., Shliaha, P. V., Miele, M., Hendrickson, R. C., Jiang, X., & Helin, K. (2023). H3K4me3 regulates RNA polymerase II promoter-proximal pause-release. Nature, 615(7951), 339–348. 10.1038/s41586-023-05780-8

Wang, H., & Helin, K. (2024). Roles of H3K4 methylation in biology and disease. In Trends in Cell Biology. Elsevier Ltd. 10.1016/j.tcb.2024.06.001

Wang, P., Lin, C., Smith, E. R., Guo, H., Sanderson, B. W., Wu, M., Gogol, M., Alexander, T., Seidel, C., Wiedemann, L. M., Ge, K., Krumlauf, R., & Shilatifard, A. (2009). Global Analysis of H3K4 Methylation Defines MLL Family Member Targets and Points to a Role for MLL1-Mediated H3K4 Methylation in the Regulation of Transcriptional Initiation by RNA Polymerase II. Molecular and Cellular Biology, 29(22), 6074–6085. 10.1128/mcb.00924-09

Wang, W., Chen, Z., Mao, Z., Zhang, H., Ding, X., Chen, S., Zhang, X., Xu, R., & Zhu, B. (2011). Nucleolar protein Spindlin1 recognizes H3K4 methylation and stimulates the expression of rRNA genes. EMBO Reports, 12(11), 1160–1166. 10.1038/embor.2011.184

Werner, F., & Grohmann, D. (2011). Evolution of multisubunit RNA polymerases in the three domains of life. In Nature Reviews Microbiology (Vol. 9, Issue 2, pp. 85–98). 10.1038/nrmicro2507

Wu, M., Wang, P. F., Lee, J. S., Martin-Brown, S., Florens, L., Washburn, M., & Shilatifard, A. (2008). Molecular Regulation of H3K4 Trimethylation by Wdr82, a Component of Human Set1/COMPASS. Molecular and Cellular Biology, 28(24), 7337–7344. 10.1128/mcb.00976-08

Wu, M., Wei, W., Chen, J., Cong, R., Shi, T., Li, J., Wong, J., & Du, J. X. (2017). Acidic domains differentially read histone H3 lysine 4 methylation status and are widely present in chromatin-associated proteins. Science China Life Sciences, 60(2), 138–151. 10.1007/s11427-016-0413-3

Yano, T., Nakamura, T., Blechman, J., Sorio, C., Dang, C. V, Geiger, B., & Canaani, E. (1997). Nuclear punctate distribution of ALL-1 is conferred by distinct elements at the N terminus of the protein (specklestrithorax11q23 abnormalities). In Cell Biology (Vol. 94). www.pnas.org.

Yu, F., Shen, X., Fan, L., & Yu, Z. (2015). Analysis of histone modifications at human ribosomal DNA in liver cancer cell. Scientific Reports, 5. 10.1038/srep18100

Yu, X., Yang, L., Li, J., Li, W., Li, D., Wang, R., Wu, K., Chen, W., Zhang, Y., Qiu, Z., & Zhou, W. (2019). De Novo and Inherited SETD1A Variants in Early-onset Epilepsy. Neuroscience Bulletin, 35(6), 1045– 1057. 10.1007/s12264-019-00400-w

Zargar, Zaffer Ullah, Mallikharjuna Rao Kimidi, and Shweta Tyagi. 2018. “Dynamic Site-Specific Recruitment of RBP2 by Pocket Protein P130 Modulates H3K4 Methylation on E2F-Responsive Promoters.” Nucleic acids research 46(1): 174–88. http://www.ncbi.nlm.nih.gov/pubmed/29059406.

Zentner, G. E., Saiakhova, A., Manaenkov, P., Adams, M. D., & Scacheri, P. C. (2011). Integrative genomic analysis of human ribosomal DNA. Nucleic Acids Research, 39(12), 4949–4960. 10.1093/nar/gkq1326

Zhang, Y., Sikes, M. L., Beyer, A. L., & Schneider, D. A. (2008). *The Paf1 complex is required for efficient transcription elongation by RNA polymerase I*. www.pnas.orgcgidoi10.1073pnas.0812939106

