## Supplemental Information for "H3K4me3 methyltransferase KMT2F promotes pre-initiation complex formation by RNA Polymerase I to regulate ribosomal RNA transcription"

**Supplementary Figure Legends**

**Supplementary Figure 1: Endogenous KMT2A and KMT2F localize to nucleolus.**

**A-B**. The immunoblots and the immunofluorescence staining (IFS) of cells either treated with control, KMT2A (**A**) or KMT2F (**B**) siRNAs are shown. Upper panel: Blots were probed with KMT2A (**A**) or KMT2F (**B**) and α-tubulin antibody as shown. Lower panel represents the loss of nucleolar localization of KMT2A (**A**) and KMT2F (**B**) upon RNAi. After siRNA treatment, cells were fixed and co-stained with either KMT2A or KMT2F and B23 antibodies. DNA was stained with 4, 6-diamidino-2-phenylindole (DAPI, blue). Scale bar, 10 µm.

**C-D**. Nucleolar localization of endogenous KMT2A (**C**) and KMT2F (**D**) is shown in three different cell lines viz: IMR90-tert, HeLa and MCF-7. Cells were co-stained with KMT2A or KMT2F and B23 (DAPI, blue). Scale bar, 10 µm.

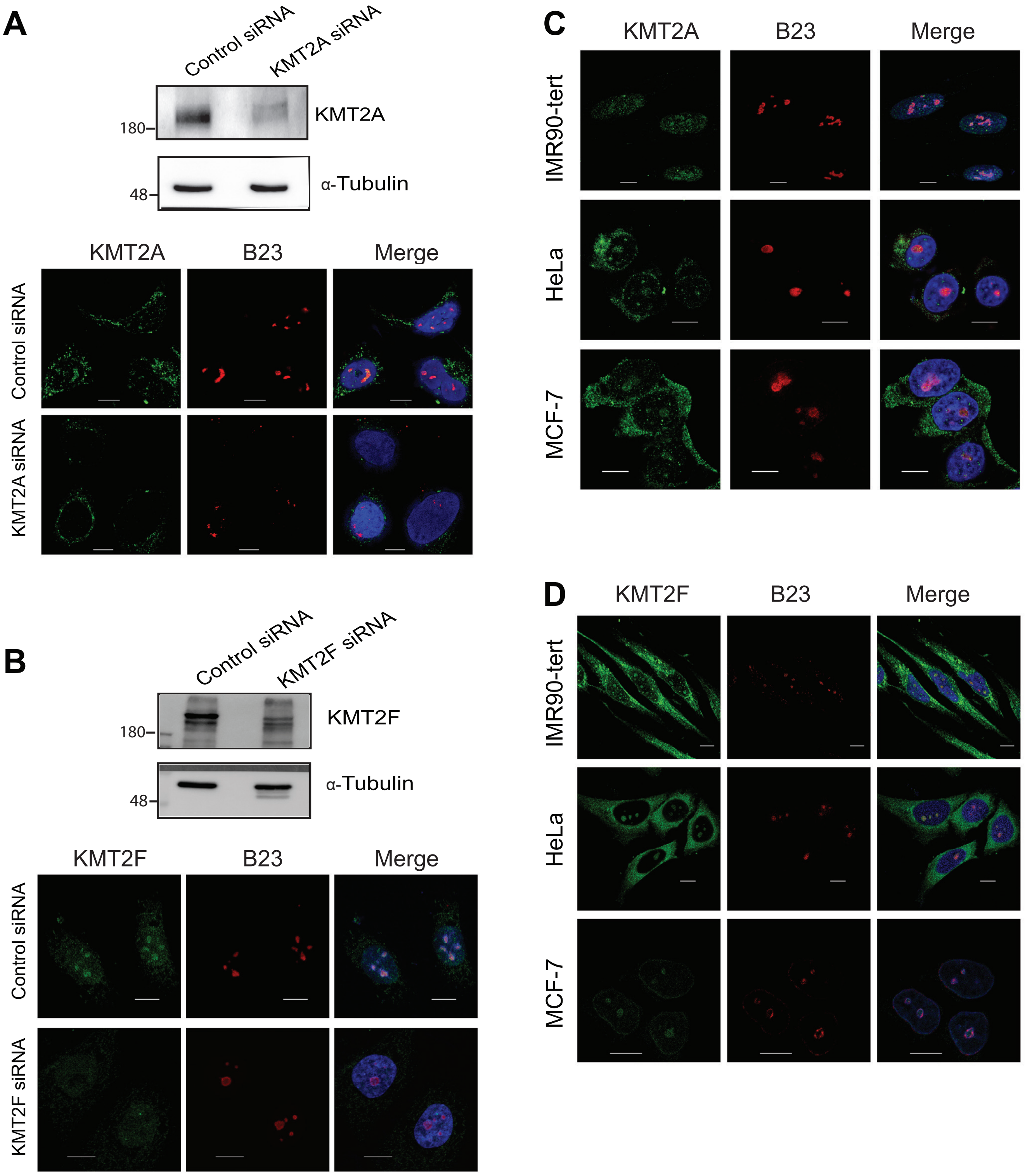


Lone Supplementary Figure 1

**Supplementary Figure 2: Epigenetic signature of active human rDNA.**

**A.** Chromatin immunoprecipitation-sequencing (ChIP-seq) maps illustrating the binding of RNA Pol I (RPA194), UBF, WDR82, WDR5, KMT2A, KMT2F, H3K4me2 and H3K4me3 at the entire human rDNA locus are shown. ChIP enrichment for each factor is expressed as the ratio of immunoprecipitated ChIP-seq signal to Input DNA ChIP-seq signal. The vertical scale (log2) in each panel represents the enrichment of each component relative to the input DNA dataset. In the assembly annotation, Subin et al. (Subin et al., 2023) excised 9.18 kb from the end of the rDNA repeat and attached it to the beginning of the repeat. As a result, the 47S promoter lies at 9.18 kb from start. Therefore, the scale shows — 9.18 to 34.8 kb with 47S promoter denoted as 0. Inset of dashed lines is shown in Figure 1B and 2A.

ChIP-seq signals

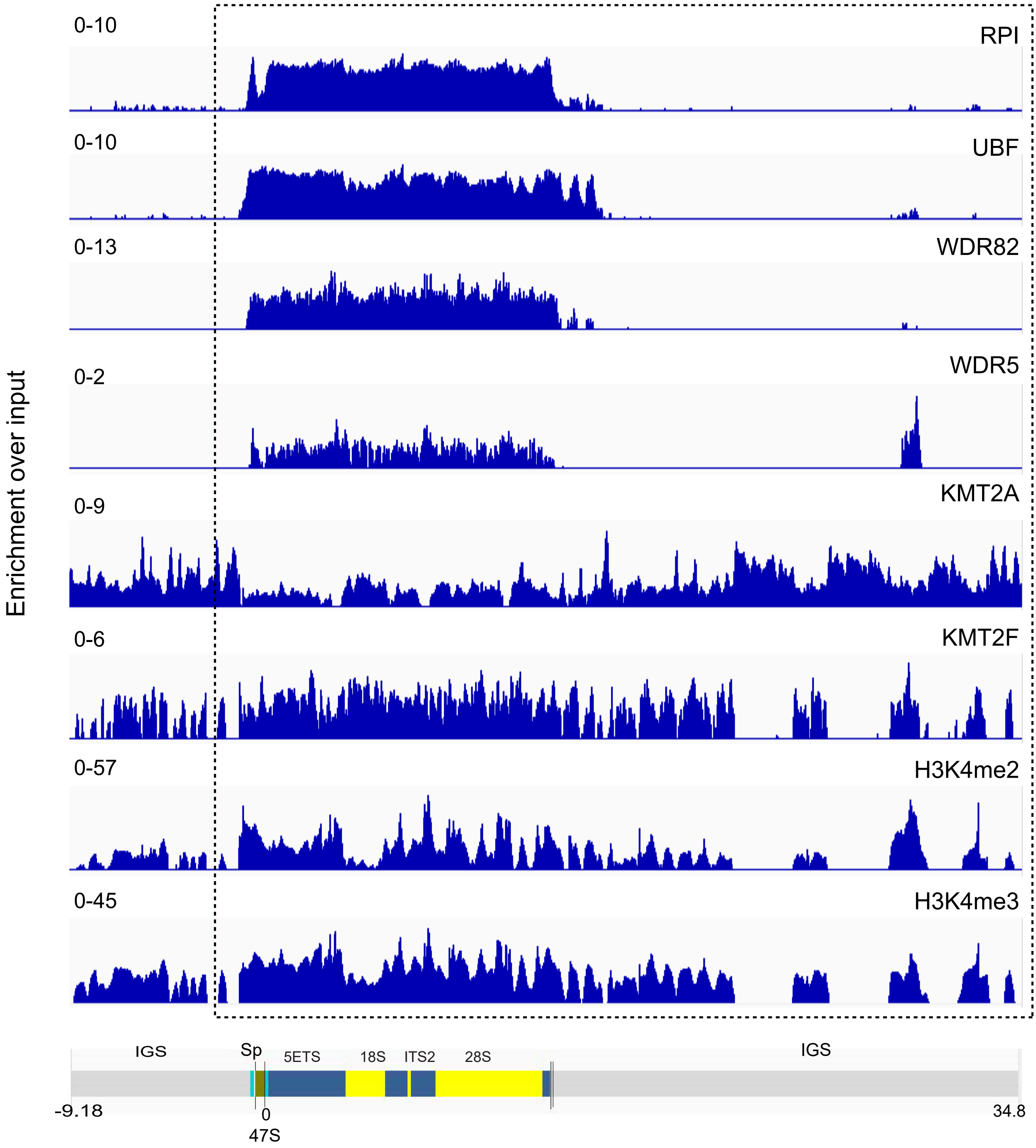


Lone Supplementary Figure 2

**Supplementary Figure 3. Binding of KMT2 and RNA Pol I to human rDNA repeats**

**A-E**. ChIP of RNA Pol I (**A**), UBF (**B**), KMT2A (**C**), KMT2F (**D**), H3(**E**) followed by qRT-PCR analysis is shown. KMT2A and KMT2F ChIP was done in IMR-90 tert cells, whereas RNA Pol I, UBF and H3 ChIP was done in HEK-293 cells. Primers 1-18 were used to assess the binding of KMT2A/KMT2F over the 43kb human rDNA repeat. Spacer promoter, terminator element (T0) and 47S (core) promoter are represented by primers 1- 3; transcribed region: 4-8 and IGS by 9-18. HOXA9 and RAD18 were used as positive control regions for KMT2A and KMT2F binding respectively while CD4 was used as negative control. RNA Pol I and UBF binding was analyzed in promoter and in the coding region of rDNA (primer 1 to 8, primer 9 as control). Data is represented from three or more than three experiments. Error bars represent SD. *$P \leq 0.05$, ** $P \leq 0.005$, ***$P \leq 0.0005$, ****$P \leq 0.00005$, ns: not significant $P > 0.05$ (two- tailed Students *t* test). Green: RNA Pol I promoter; blue: RNA Pol I transcribed region; orange: IGS.

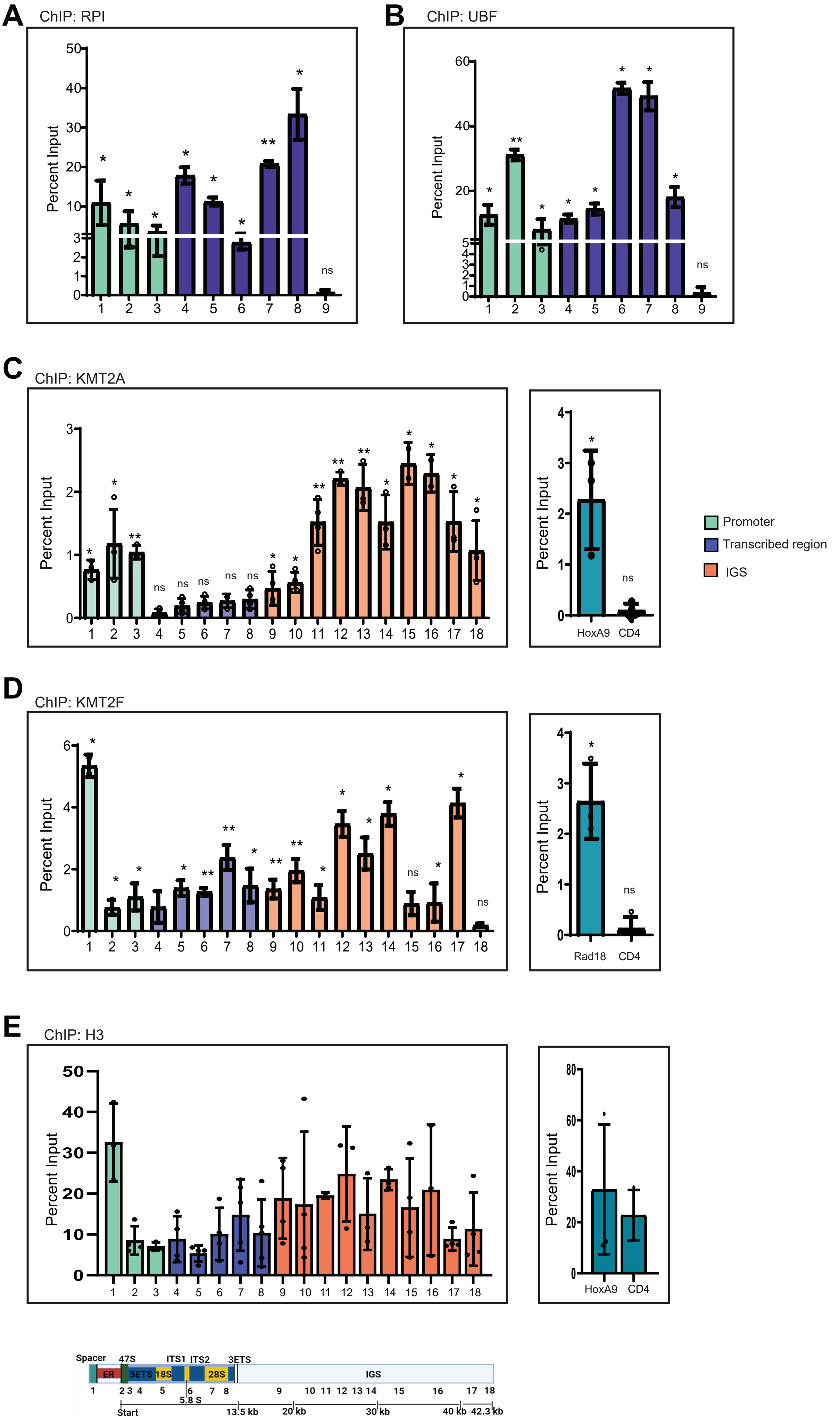


Lone Supplementary Figure 3

**Supplementary Figure 4: RNAi mediated depletion of KMTs affect ribosomal DNA transcription.**

**A.** HeLa cells were subjected to endogenous immunoprecipitation (IP) using antibody against the N terminal subunit of KMT2A. The anti-immunoglobulin G (IgG)-mock antibody was used as control. The immunoblot was probed with α-RPA194, α-UBF, α- RRN3 and α-KMT2A antibodies. The molecular weight marker is indicated on the left.

**B-D**. qRT-PCR analysis of KMT2A (**A**), KMT2F (**B**) and WDR5 (**C**) transcript levels after treating the cells with Control, KMT2A, KMT2F and WDR5 siRNA is shown. Error bars denote SD. $*P \leq 0.05$, $** P \leq 0.005$, $***P \leq 0.0005$, ns: not significant $P > 0.05$ (two-tailed Students *t* test).

**E.** Schematic of KMT2A and KMT2A mutant used in transcriptional analysis in Figure 4 are shown. Recombinant full-length KMT2A (FL) was generated by PCR sub-cloning. KMT2A mutant devoid of Su(var)3-9, Enhancer-of-zeste and Trithorax (SET; KMT2AΔSET) and Transactivation domain (TAD; KMT2AΔ TAD) were generated by deletion PCR.

**F**. Shows qRT-PCR analysis of KMT2A transcript levels in control cells, KMT2A full length (2A-FL), KMT2AΔSET (2AΔSET), and KMT2AΔTAD (2AΔTAD) cells.. Error bars denote SD. $*P \leq 0.05$, $** P \leq 0.005$, $***P \leq 0.0005$, ns: not significant $P > 0.05$ (two-tailed Students *t* test)

**G.** Schematic of KMT2F full length construct and KMT2F mutant (KMT2FΔSET) are shown. siRNA resistant (SR) KMT2F construct was generated by PCR based mutagenesis resulting in silent mutations. Subsequently, KMT2F mutant lacking SET domain was generated by deletion PCR from FL SR construct.

**H**. qRT-PCR analysis of KMT2F transcript levels in control cells, KMT2F full length (2F-FL), and KMT2FΔSET (2FΔSET) cells. Error bars denote SD. *P ≤ 0.05, ** P ≤ 0.005, ***P≤ 0.0005, ns: not significant P > 0.05 (two-tailed Students *t* test)

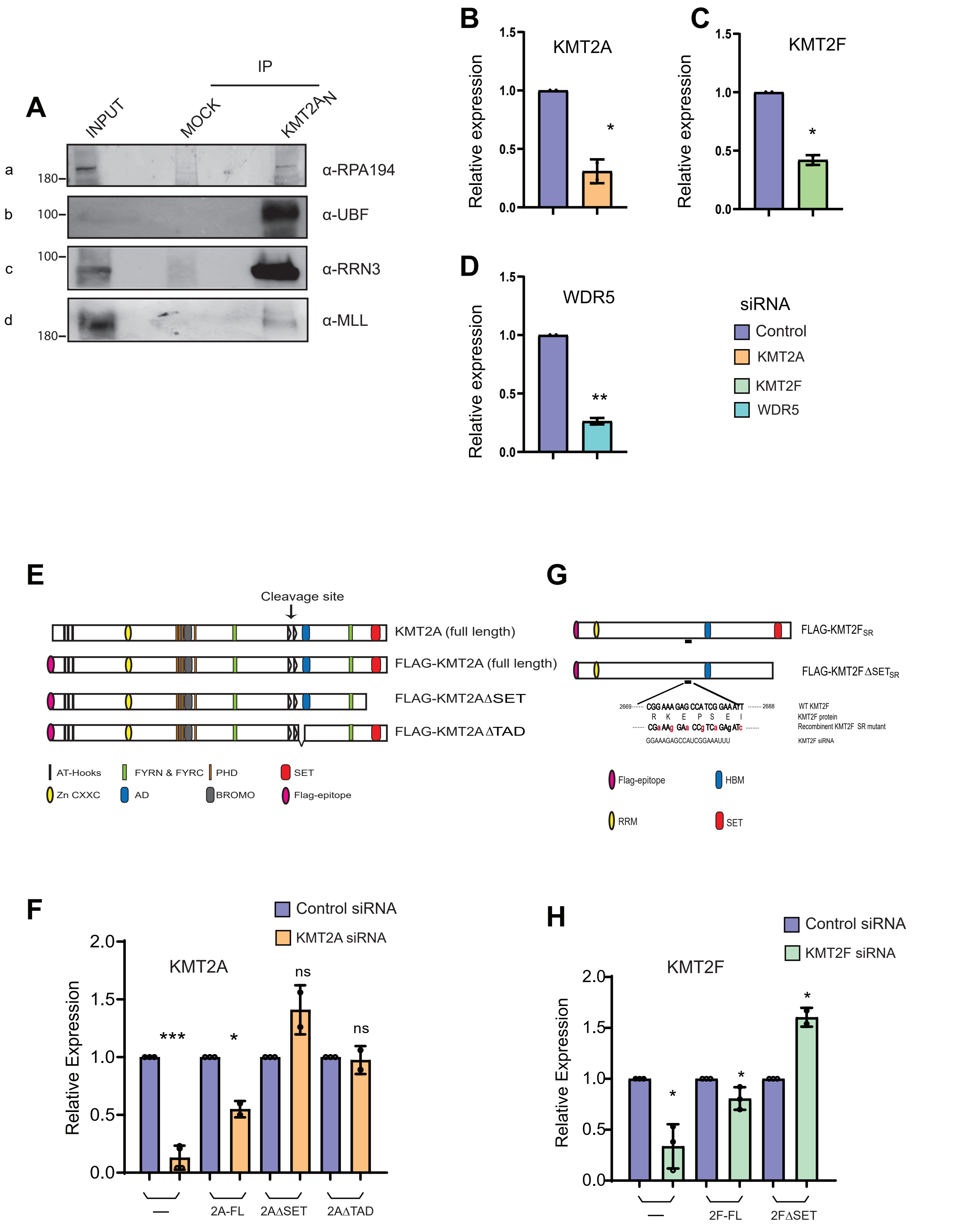


Lone Supplementary Figure 4

**Supplementary Figure 5: Depletion of KMTs affect ribosomal DNA transcription in different cell lines.**

**A.** Representative autoradiogram displaying the $^{32}P$ orthophosphate-labeled ribosomal RNA (45S and 32S) following KMT2A loss in various KMT2A mutant cell lines. U-2OS cells were treated with either control or KMT2A siRNA. Likewise, KMT2A F.L, KMT2AΔSET and KMT2AΔTAD cells received the same treatments. The total RNA extracted was analyzed on an agarose gel and visualized through autoradiography (upper panel). Additionally, 28S RNA levels were assessed using ethidium bromide staining on a formaldehyde agarose gel (lower panel).

**B.** Representative autoradiogram showing the $^{32}P$ orthophosphate-labeled ribosomal RNA (45S and 32S) after KMT2F loss in various KMT2F mutant cell lines. U-2OS cells were treated with either control or KMT2F siRNA. Similarly, KMT2F F.L and KMT2FΔSET cells received the same treatments and were subjected to RNA extraction. The extracted total RNA was analyzed on an agarose gel and visualized via autoradiography (upper panel). Additionally, 28S RNA levels were evaluated using ethidium bromide staining on a formaldehyde agarose gel (lower panel).

**C.** Represents the autoradiogram depicting the $^{32}P$ orthophosphate-labeled ribosomal RNA (45S and 32S) after the loss of different KMT2 members (KMT2A, KMT2F, KMT2B, KMT2C) and WDR5.

**D.** Effect of KMT2A loss on rRNA transcription in different cell lines is shown. HeLa, MCF7 and IMR90-tert cells were treated with KMT2A siRNA or control siRNA. Results here show densitometry quantification of $^{32}P$ incorporation into 45S ribosomal RNA levels upon KMT2A loss. Each value represents the result of two independent experiments, with standard deviation indicated by error bars. Statistical significance was assessed using a two-tailed Student's *t* test: $*P \leq 0.05$, $**P \leq 0.005$,.

**E.** Effect of KMT2F loss on ribosomal transcript levels in HeLa and MCF7 cells. Cells were treated with KMT2F or Control siRNA and subjected to densitometry quantification of $^{32}P$ incorporation into 45S ribosomal RNA levels. The relative intensity was measured by normalizing 45S levels with 28S rRNA loading control. Each value is an outcome of two independent experiments. S.D is represented by error bars. The statistical significance was calculated by two-tailed student *t* test $**P \leq 0.005$.

**F.** IFS of endogenous KMTB is shown in U-2OS cells. Cells were co-stained with KMT2B and B23 antibodies (DAPI, blue). Scale bar, 10 µm.

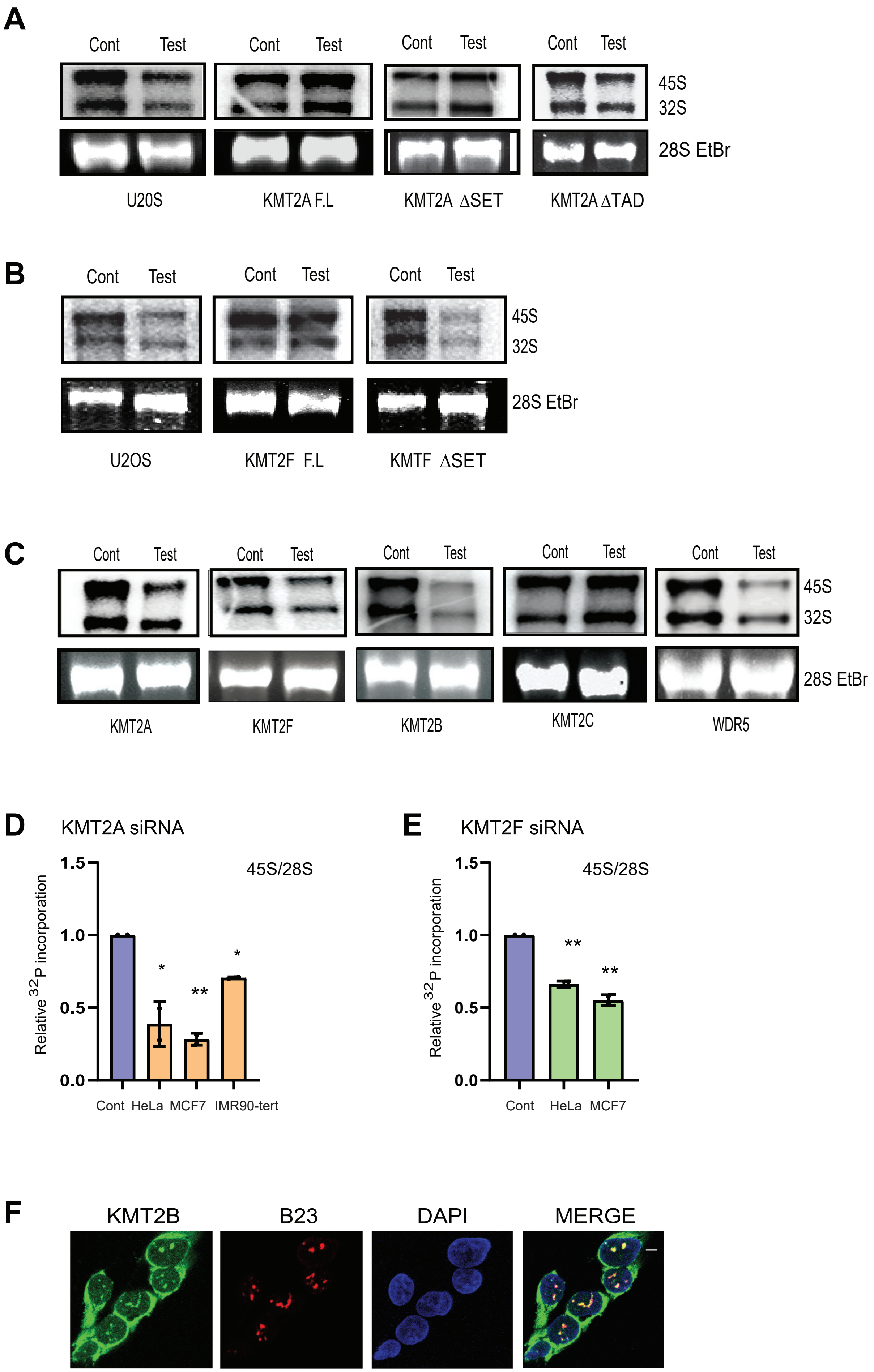


Lone Supplementary Figure 5

**Supplementary Figure. 6. KMT2F deposits H3K4me3 marks to regulate epigenetic state of human rDNA.**

**A-C**. Represents ChIP analysis of KMT2F (**A**), H3K4Me2(**B**), H3K4Me3 (**C**) in KMT2F shRNA KD. RAD18 and CD4 were used as positive and negative control primers. (A-C) Experiments were performed three or more than three times as biological replicates. Error bars represent SD. $*P \leq 0.05$, $**\ P \leq 0.005$, $***P \leq 0.0005$, $****P \leq 0.00005$, ns: not significant $P > 0.05$ (two- tailed Students *t* test)

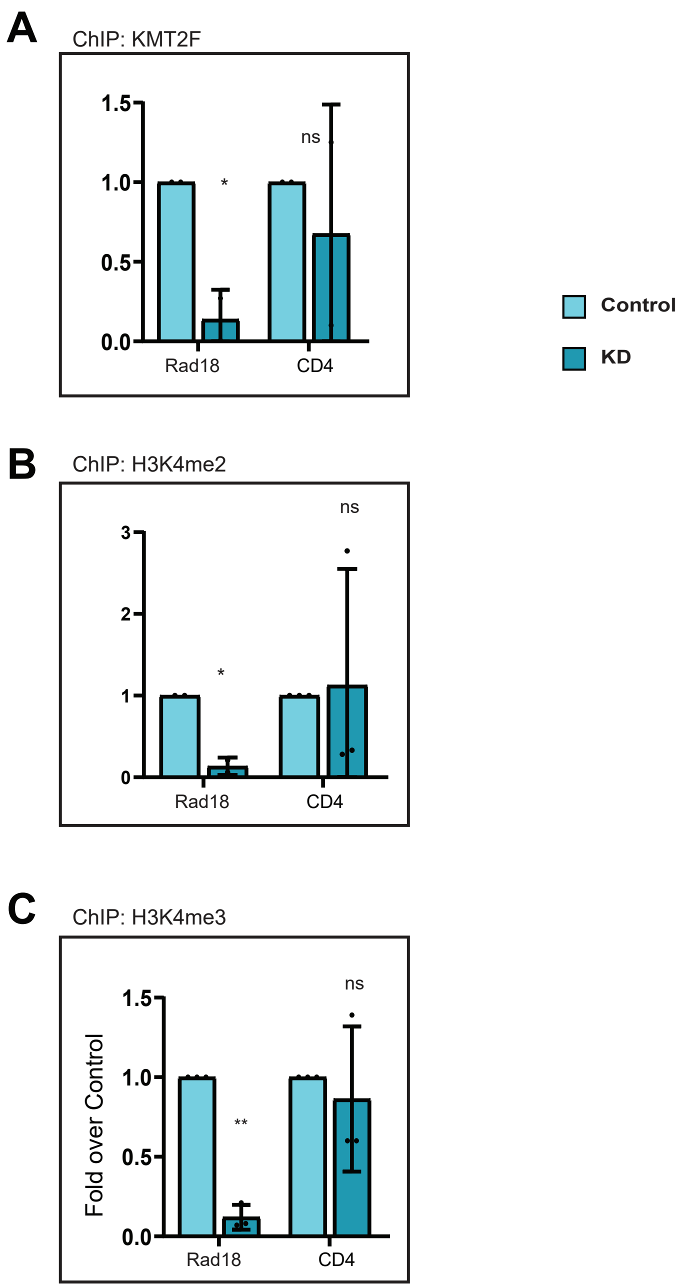


Lone Supplementary Figure 6

**Supplementary Figure. 7. Loss of KMT2F affect the formation of RNA Pol I pre-initiation complex on rDNA.**

**A-B**. ChIP analysis of H3K9ac (**A**) and H3K9me3 (**A**) in KMT2F shRNA KD. RAD18 and CD4 were used as positive and negative control primers. Error bars indicate standard deviation (SD). Significance levels are as follows: *P ≤ 0.05, **P ≤ 0.005, ***P ≤ 0.0005, ****P ≤ 0.00005, ns: not significant (P > 0.05, two-tailed Student's *t* test).

**C.** A schematic of the full-length KMT2F construct, along with its various domains, was used to PCR amplify the RRM domain fragment spanning amino acids 84 to 172. This fragment was then cloned into a vector containing the SFB tag. Ectopically expressed SFB-RRM and SFB were subjected to S-protein pulldowns. The RRM pull-down was detected using anti-FLAG antibody, and the immunoblot was probed with various antibodies (RPA194, RRN3, and RPA-49) to assess the interaction of RRM with different components of the pre-initiation complex. SFB, used as a mock control, was also detected with the anti-FLAG antibody.

**D**. The full-length KMT2F (2F wt) and the KMT2F mutant (2F mut), along with SFB (mock) were used to study the interaction of the KMT2F in the wild type and in the mutant conditions with the components of the pre-initiation complex. These constructs were overexpressed in cells and subjected to S-protein pulldowns using S protein beads. The pull-downs of both KMT2F wt and KMT2F mut were detected using anti-FLAG antibody. Immunoblots were probed with various antibodies for the pre-initiation complex (RPA-194, RRN3, and RPA-49) to assess the interaction of KMT2F in both wild-type and mutant conditions.

**E.** Represents the quantification of protein interaction levels between KMT2F and the various components of the pre-initiation complex (RPA-194, RRN3 and RPA-49) under wild-type and mutant conditions, as shown in **D**. The quantification was performed by normalizing the interactions in mutant conditions to those in wild-type conditions. Error bars indicate standard deviation (SD). ns indicates not significant P > 0.05, two-tailed Student's *t* test.

**F.** Represents the ChIP analysis of the RPA-194 (RNA Pol I) and UBF done in the control cells or in the stable cell lines of the KMT2F expressing KMT2F Full length represented as "wt" or in the catalytically dead KMT2F represented as "mut", both the cell lines were treated with 3UTR shRNA targeting the degradation of the endogenous KMT2F. ChIPed DNA in these three conditions was subjected to qRT-PCR and the binding of the RPI and UBF rDNA promoter was detected. Significance was calculated with respect to the control in both the wt and mutant conditions using two-tailed Student's *t* test. Error bars indicate standard deviation (SD), *P ≤ 0.05, **P ≤ 0.005, and ns indicates not significant (P > 0.05)

**G.** The expression levels of KMT2A (2A) and KMT2F (2F) in overexpressed KMT2A and KMT2F HEK293 cells is shown. Data is presented from three or more than three experiments. Significance is calculated with respect to the control cells, or in between the Error bars represent SD. *P ≤ 0.05, ** P ≤ 0.005, ***P≤ 0.0005, ****P≤ 0.00005, ns: not significant P > 0.05 (two-tailed Students t test). ND, not done.

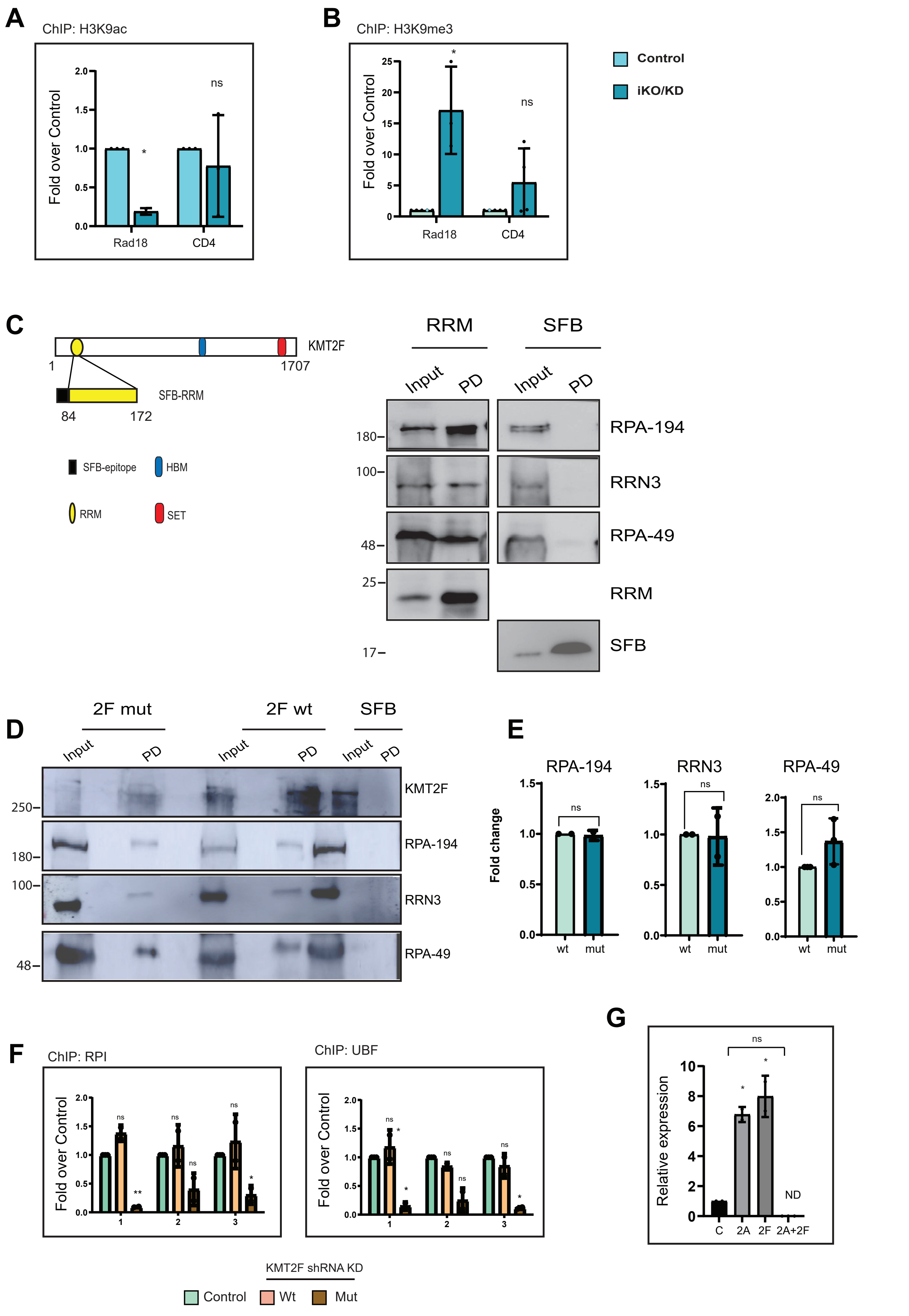

Lone Supplementary Figure 7

**Supplementary Table S1: ChIP and transcript primers used in this study.**

Primer sequences, with their coordinates on rDNA are provided here.
